# Paralogous guanine deaminases acquired from bacteria by horizontal gene transfer promote purine homeostasis in *Caenorhabditis elegans*

**DOI:** 10.64898/2026.04.09.715363

**Authors:** Sushila Bhattacharya, Lisa Fischer, Evrim Fer, Jennifer Snoozy, Grace N. Hagedorn, Marco Herde, Betül Kaçar, Claus-Peter Witte, Kurt Warnhoff

## Abstract

Disruptions in purine metabolism contribute to a range of human diseases, from rare genetic disorders such as Lesch-Nyhan syndrome and xanthinuria to common conditions including gout and cancer. To better understand the metabolic networks that regulate purine homeostasis, we developed a *Caenorhabditis elegans* model of xanthine dehydrogenase (*xdh-1*) deficiency. Remarkably, *xdh-1* mutant animals form rare xanthine stones, recapitulating a hallmark of human xanthinuria. To uncover genetic regulators of purine homeostasis, we performed a forward genetic screen for mutations that exacerbate xanthine stone formation in *xdh-1* mutants. This approach identified multiple loss-of-function alleles in a previously uncharacterized gene, which we named *gda-1*. We show that *gda-1* encodes an intestinal guanine deaminase that mediates a key enzymatic step in purine catabolism. The *C. elegans* genome also encodes a paralog, *gda-2*, which shares guanine deaminase activity but is expressed in distinct tissues. While *gda-2* can compensate for *gda-1* loss in guanine metabolism, the two genes exhibit non-redundant roles in regulating xanthine accumulation and stone formation. Interestingly, our evolutionary analyses suggest that *gda-2* was acquired by nematodes via horizontal gene transfer from bacteria. These findings reveal a spatially regulated purine catabolism pathway in *C. elegans* and suggest that acquisition of bacterial genes has shaped a core nematode metabolic network.

## Introduction

Life depends on intricate metabolic networks that not only sustain cellular function but also evolve in response to shifting environments and ecological pressures. These networks are dynamic systems, shaped by evolutionary forces such as gene duplication, divergence, and, in some cases, horizontal gene transfer (HGT) [1, 2]. Such evolutionary processes enable organisms to adapt to changing nutrient landscapes, yet the extent to which HGT contributes to core metabolic pathways in animals remains poorly understood.

Horizontal gene transfer, the movement of genetic material between unrelated organisms, contrasts with the traditional vertical inheritance of genes from parent to offspring. While HGT is common among prokaryotes and underlies phenomena such as the spread of antibiotic resistance [3], its occurrence in animals is considered rare. Nevertheless, striking examples reveal that HGT can profoundly impact animal biology: it has enabled beetles to exploit novel nutrient sources [4], empowered predator evasion in aphids [5], and enhanced stress tolerance, plant parasitism, and nutrient acquisition in nematodes [6–9], among other cases. These examples represent evolutionary leaps, where organisms have acquired biochemical solutions from entirely different domains of life. These rare but transformative events raise fascinating questions about how such genes integrate into existing networks.

Purine metabolism provides a useful context to explore these questions. This pathway governs nucleotide turnover and energy balance, and its disruption underlies a spectrum of human disorders, from rare genetic diseases such as Lesch-Nyhan syndrome and xanthinuria to common conditions including gout and cancer [10–13]. The purine catabolic pathway in humans culminates in the oxidation of xanthine to uric acid by xanthine dehydrogenase (XDH, **Fig. 1A**) [14]. Loss of XDH activity in humans causes xanthinuria, characterized by excessive xanthine accumulation, stone formation, and sometimes renal failure [15, 16]. Although the enzymology of purine catabolism is well described, less is known about how these steps are integrated across tissues and how organisms compensate when key reactions are disrupted. Model organisms offer powerful systems to address these gaps.

**Figure 1.**
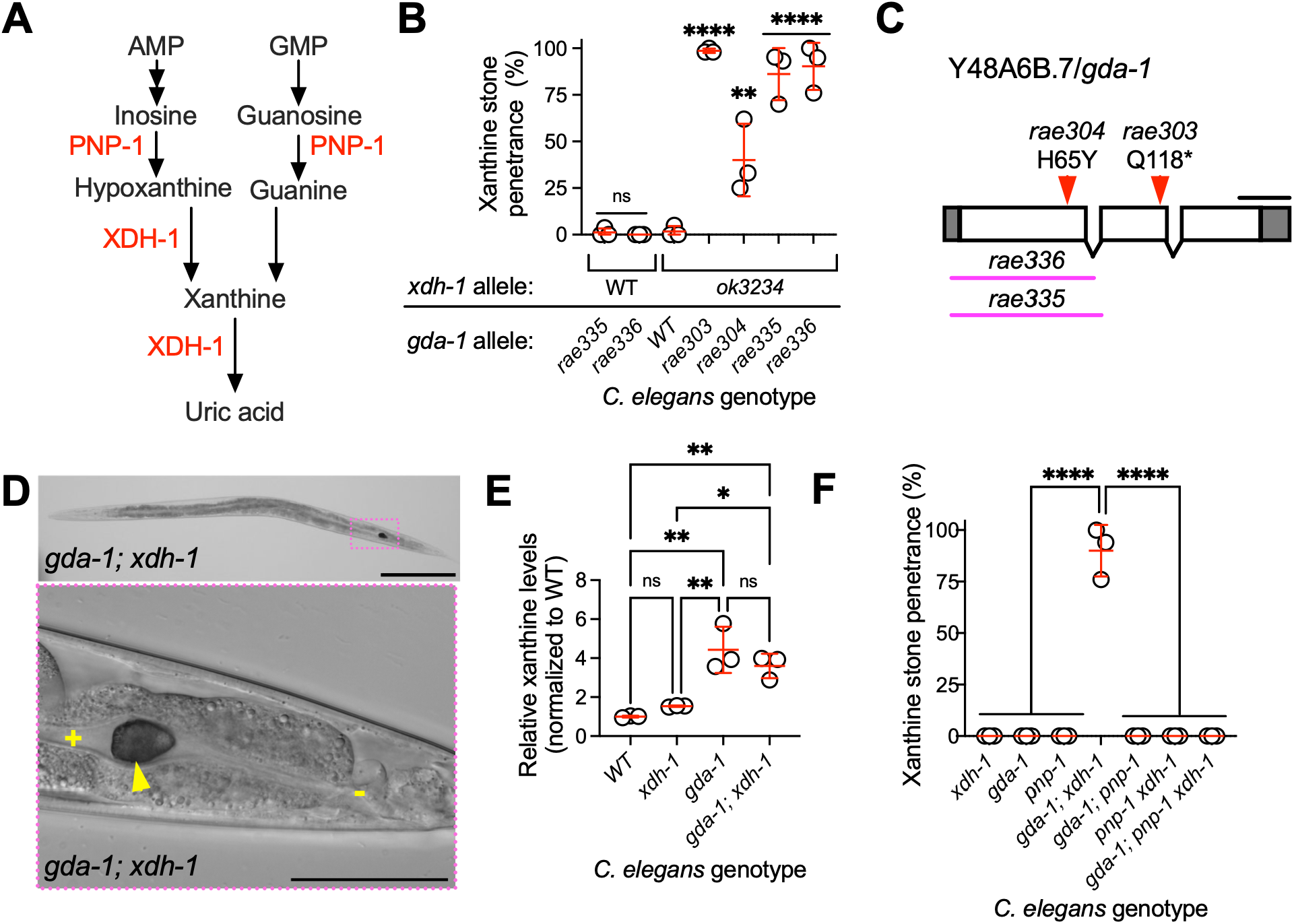
*gda-1*/Y48A6B.7 inhibited the accumulation of xanthine. A) Simplified schematic of purine catabolism in *C. elegans*, highlighting the roles of purine nucleoside phosphorylase (PNP-1) and xanthine dehydrogenase (XDH-1). B) Xanthine stone formation in *C. elegans* carrying various *gda-1* mutant alleles (*rae303*, *rae304*, *rae335*, *rae336*), *xdh-1(ok3234)*, and double mutants combining *xdh-1* with *gda-1* alleles. Animals were assayed for xanthine stone formation. Data points represent biological replicates. Mean and standard deviation are displayed. ns, not significant; **, p<0.01; ****, p<0.0001; ordinary one-way ANOVA. C) Structure of the *gda-1* gene. Boxes represent exons; connecting lines indicate introns. *rae303* and *rae304* are chemically induced alleles identified in a screen for mutations that enhance xanthine stone formation in an *xdh-1(ok3234)* background. *rae335* and *rae336* are CRISPR/Cas9-generated deletions. Scale bar: 100 bp. D) Differential interference contrast images showing a xanthine stone in the posterior intestine of a *gda-1; xdh-1* young adult *C. elegans*. The arrowhead marks the xanthine stone; “+” indicates the intestinal lumen; “-” indicates the rectum. Scale bars: 100 μm (upper) and 50 μm (lower). E) Quantification of xanthine in wild-type, *xdh-1, gda-1,* and *gda-1; xdh-1* mutant young adult *C. elegans*. Each data point represents a biological replicate, and values are normalized to the wild type. ns, not significant; *, p<0.05; **, p<0.01; ordinary one-way ANOVA. Note, these data are derived from the same experiment displayed in **Fig. S8**. F) Xanthine stone formation in various mutant *C. elegans* was assessed. Data points represent biological replicates. Mean and standard deviation are displayed. ****, p<0.0001; ordinary one-way ANOVA. Sample size and individuals scored per biological replicate are provided in **Table S1**.

Here, we leverage the genetic tractability of the nematode *Caenorhabditis elegans* to dissect purine homeostasis *in vivo*. Using an *xdh-1* mutant that recapitulates a hallmark of human xanthinuria, xanthine stone formation, we performed a genetic screen to identify modifiers of purine metabolism [17]. This approach revealed loss-of-function mutations in *gda-1*, an intestinal guanine deaminase, and prompted investigation of its highly similar paralog, *gda-2*. Although both enzymes share guanine deaminase activity, they exhibit distinct tissue expression and non-redundant roles in purine flux. Strikingly, phylogenetic analyses suggest that *gda-2* originated via horizontal gene transfer from bacteria, revealing that nematodes have incorporated a microbial enzyme into a core metabolic network. Together, these results reveal an unexpected role for horizontal gene transfer in promoting metabolic robustness in an animal.

## Results

### *gda-1/Y48A6B.7* inhibited the accumulation of xanthine

To better understand mechanisms of purine homeostasis, we previously established a *C. elegans* model of human xanthinuria, a rare disease caused by inactivation of xanthine dehydrogenase and characterized by elevated levels of xanthine in blood and urine, accumulation of xanthine kidney stones, and renal failure [15, 17]. *C. elegans* with a null mutation in the gene encoding for xanthine dehydrogenase, *xdh-1,* develop highly autofluorescent xanthine stones. However, these stones are rare, only developing in about 2% of *xdh-1* mutant animals [17]. This result is surprising given the central role of XDH-1 in the catabolism of purines and suggests alternate mechanisms for the maintenance of purine homeostasis (**Fig. 1A**).

To identify additional genetic regulators of purine homeostasis, we performed an unbiased genetic screen for mutations that enhanced the penetrance of xanthine stone formation in *xdh-1* mutant animals. We mutagenized *xdh-1* mutant *C. elegans* with ethyl methane sulfonate (EMS) and cultured the newly mutagenized animals for two generations allowing newly induced mutations to become homozygous [18]. Individual mutagenized F2 animals were isolated and screened to identify strains in which a high fraction of F3 progeny developed xanthine stones. This screen previously revealed five loss-of-function mutations in *sulp-4,* a sulfate permease that negatively regulates xanthine stone accumulation in parallel to the activity of *xdh-1* [17].

Here we describe two new EMS-induced recessive loss-of-function mutations, *rae303* and *rae304,* that caused a high penetrance of xanthine stone formation in the posterior lumen of the *C. elegans* intestine when combined with an *xdh-1* mutation (**Fig. 1B-D**). These mutations were prioritized because they displayed strong enhancement of xanthine stone formation and formed a complementation group, indicating they affect a single gene (see **Materials and Methods)**. To identify the causative genetic lesions in these new mutant strains, genomic DNA was isolated and analyzed via whole genome sequencing. Given that the *rae303* and *rae304* mutations fail to complement one another, we predicted that these mutant strains should each have a novel EMS-induced mutation in a common gene. Upon bioinformatic analysis of our whole genome sequencing, only one gene was uniquely impacted in both mutant strains, *gda-1*/Y48A6B.7, strongly suggesting that these mutations in *gda-1* were causative for the enhanced penetrance of xanthine stone formation in *xdh-1* mutant animals (**Fig. 1C**). *rae303* was a nonsense allele of *gda-1* that encoded a premature stop codon in place of glutamine at amino acid 118 (chromosome III, position 11025882, G>A, Q118*). *rae304* was a missense allele which encoded a tyrosine in place of a histidine residue at amino acid position 65 (chromosome III, position 11026349, G>A, H65Y, **Fig.1C**). Given that *rae303* and *rae304* were recessive mutations and *rae303* caused a premature stop codon, we propose that *rae303* and *rae304* encode strong loss-of-function or null alleles of *gda-1*.

To test the hypothesis that loss of *gda-1* caused enhanced xanthine stone formation in an *xdh-1* mutant background, we used CRISPR/Cas9 to engineer new *gda-1* deletion alleles, *rae335* and *rae336* [19, 20] (**Fig. 1C**). The *gda-1(rae335)* and *gda-1(rae336)* alleles were 531 bp deletion (with a 38 bp insertion) and 556 bp deletions respectively that eliminated the *gda-1* start codon as well as all of exon one. Thus, we propose that *rae335* and *rae336* encode null alleles of *gda-1.* Both *gda-1(rae335)* and *gda-1(rae336)* strongly enhanced the penetrance of xanthine stone formation caused by *xdh-1* inactivation, phenocopying the *gda-1* alleles isolated from our forward genetic screen (**Fig. 1B).** We conclude that *gda-1* is necessary for preventing the accumulation of xanthine stones in *C. elegans* also carrying an *xdh-1* mutation. Interestingly, the *gda-1* or *xdh-1* mutations do not induce substantial xanthine stone formation on their own, demonstrating that *gda-1* and *xdh-1* act in parallel genetic pathways to limit xanthine stone formation (**Fig. 1B**). We use *rae336* as our *gda-1* reference allele.

We wanted to understand if the dramatic enhancement of xanthine stone formation caused by *gda-1* inactivation was reflective of a parallel increase in whole-animal xanthine content. To determine the impact of *gda-1* inactivation on xanthine levels, we employed HPLC-MS to measure xanthine in *C. elegans* extracts. We measured xanthine in wild-type, *xdh-1, gda-1,* and *gda-1; xdh-1* mutant young-adult animals. Surprisingly, *xdh-1* single mutant animals did not have significantly elevated xanthine compared to the wild type. However, we observed a substantial 4-fold increase in xanthine in *gda-1* single mutant *C. elegans* that was also present in the *gda-1; xdh-1* double mutant animals (**Fig. 1E)**. These data demonstrate that *gda-1* is necessary for preventing the accumulation of xanthine. Intriguingly, despite no difference in the accumulation of xanthine between *gda-1* and *gda-1; xdh-1* double mutant animals, we observed a dramatic difference in the formation of xanthine stones between these two strains **(Fig. 1B,E)**. This suggests that there are factors contributing to the formation of xanthine stones in addition to xanthine accumulation.

To genetically dissect the nature of the xanthine stones that develop in *gda-1; xdh-1* animals, we introduced a mutation in *pnp-1* which encodes the *C. elegans* purine nucleoside phosphorylase [17, 21]. PNP-1 cleaves the nucleosides inosine or guanosine by phosphorolysis into ribose-1-phosphate and the bases hypoxanthine or guanine, metabolic steps required for xanthine formation (**Fig. 1A**). To test the model that *pnp-1* was necessary for the formation of xanthine stones in *gda-1; xdh-1* mutant animals, we generated *gda-1; pnp-1 xdh-1* triple mutant animals and assayed the formation of xanthine stones. *gda-1; pnp-1 xdh-1* triple mutant animals did not develop xanthine stones (**Fig. 1F**). These data support our model that xanthine produced downstream of PNP-1 is the critical component of the fluorescent stones we observe in our *gda-1; xdh-1* double mutant animals. Although, we cannot exclude the possibility that other PNP-1-derived metabolites (hypoxanthine or guanine) also contribute to stone formation.

*pnp-1* loss of function also suppresses the enhanced formation of xanthine stones caused by *sulp-4* inactivation [17]. *xdh-1; sulp-4* double mutant *C. elegans* develop xanthine stones while *pnp-1 xdh-1; sulp-4* triple mutants do not. Loss of sulfur amino acid catabolism also suppresses xanthine stone formation in *xdh-1; sulp-4* animals, likely by preventing excessive sulfate accumulation that drives osmotic imbalance and promotes stone development [17]. To test whether sulfur amino acid metabolism impacts xanthine stone formation in *gda-1; xdh-1* animals, we constructed *gda-1; xdh-1; cdo-1* triple mutant animals. *cdo-1* encodes *C. elegans* cysteine dioxygenase, a critical enzyme in sulfur amino acid breakdown [22]. Loss of *cdo-1* did not impact xanthine stone formation in *gda-1; xdh-1* mutant animals (**Fig. S1**). Thus, sulfur amino acid metabolism is not required for the enhanced xanthine stone formation conferred by *gda-1* inactivation. Taken together, these results suggest that *gda-1* and *sulp-4* are modulating the formation of xanthine stones through distinct pathways.

### *gda-1* encoded a guanine deaminase that acted in the intestine

*gda-1* is predicted to encode a 168 amino acid deaminase and shares 47% amino acid identity with a well-characterized guanine deaminase (GDA) from the bacteria *Bacillus subtilis* (*Bs*GDA, WP_367147857, **Fig. 2A**) [23, 24]. GDA enzymes catalyze the hydrolytic deamination of guanine, producing xanthine and ammonia. Given the reaction catalyzed by GDA enzymes, it is counterintuitive that loss of a guanine deaminase would lead to accumulation of its product, xanthine. Nevertheless, key substrate-binding and catalytic residues are conserved between *Bs*GDA and GDA-1 in *C. elegans* (**Fig. 2A**) [25]. Interestingly, the *rae304* lesion isolated in our forward genetic screen alters histidine 65, corresponding to histidine 53 in *Bs*GDA, that resides in the active site of the enzyme and coordinates the catalytically essential zinc ion (**Fig. 2A)** [25]. Thus, *rae304* (H65Y) likely disrupts zinc coordination of the GDA-1 active site, reducing or eliminating the activity of the enzyme. Based on this homology we hypothesize that the *gda-1*/Y48A6B.7 genetic locus encodes a *C. elegans* guanine deaminase.

**Figure 2:**
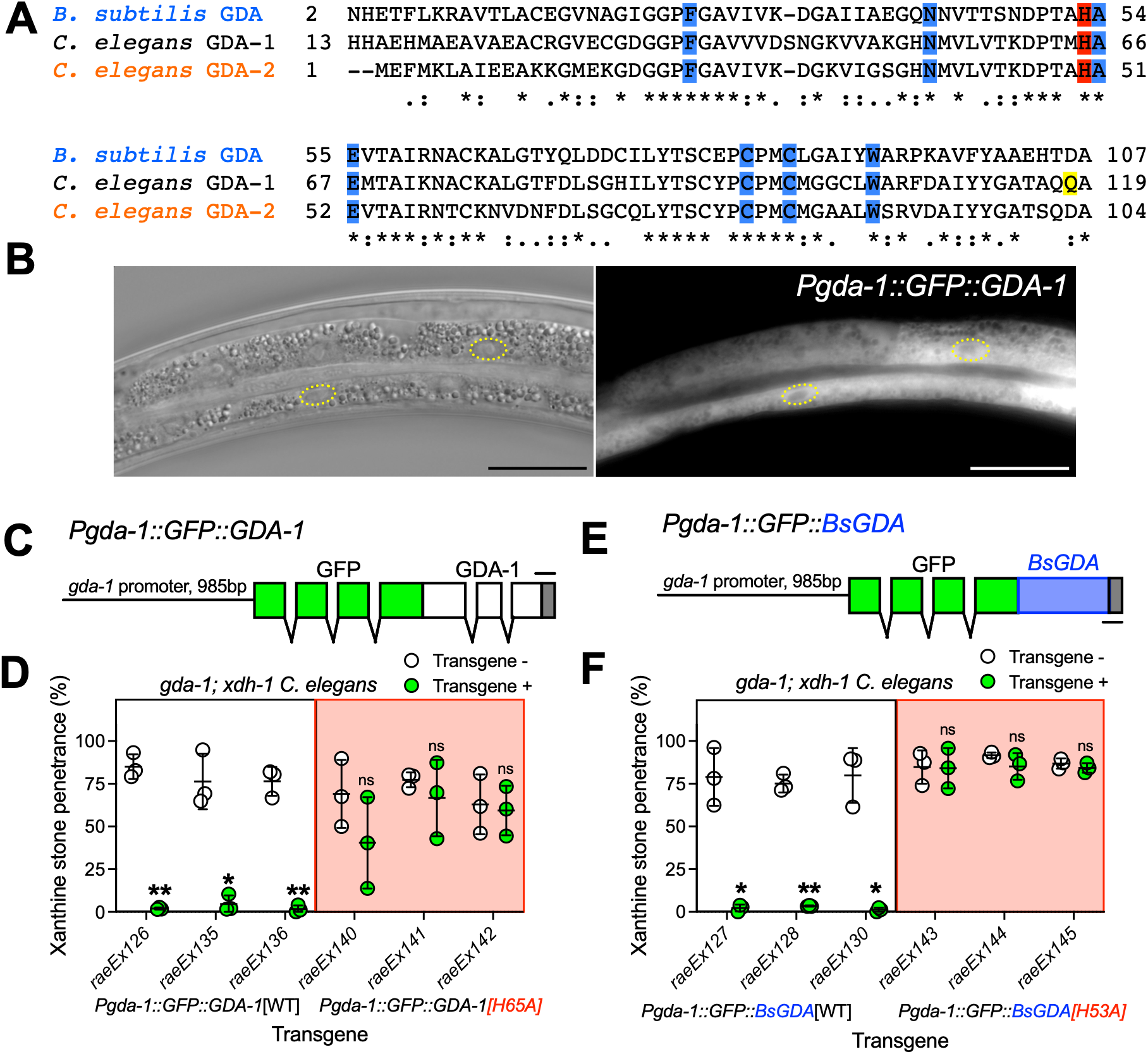
*gda-1* encoded a guanine deaminase that acted in the intestine. A) Amino acid alignment of the cytosine deaminase domain of *C. elegans* GDA-1 (NP_499418.1), GDA-2 (NP_498663.1), and *B. subtilis* GDA (WP_367147857.1). The catalytically essential histidine 65 (GDA-1) is highlighted in red. Glutamine 118 which was impacted by the *rae303* allele is highlighted in yellow. Active site and substrate binding residues are highlighted in blue. “*” indicates perfect amino acid conservation, “:” indicates strong similarity, and “.” indicates weak similarity amongst sequences compared. B) Differential interference contrast (left) and fluorescence microscopy (right) images of *gda-1; xdh-1* mutant animals expressing the *Pgda-1::GFP::GDA-1* transgene. GFP::GDA-1 localizes to the cytoplasm and nuclei (highlighted with yellow circles) of intestinal cells. Scale bars: 25 μm. C) Schematic of the *Pgda-1::GFP::GDA-1* transgene, which encodes wild-type GDA-1 fused to GFP at the N-terminus. Exons are shown as boxes, introns as connecting lines, and the 985 bp *gda-1* promoter as a straight line. Scale bar: 100 bp. D) Transgenic *gda-1; xdh-1* animals expressing either wild-type (*Pgda-1::GFP::GDA-1[WT]*) or catalytically inactive (*Pgda-1::GFP::GDA-1[H65A]*) GDA-1, along with their non-transgenic siblings, were scored for xanthine stone formation. Data points represent biological replicates. Mean and standard deviation are displayed. ns, not significant; *, p<0.05; **, p<0.01; *t* test. E) Schematic of the *Pgda-1::GFP::BsGDA* transgene, which encodes wild-type *B. subtilis* GDA fused to GFP at the N-terminus. Exons are shown as boxes, introns as connecting lines, and the 985 bp *gda-1* promoter as a straight line. Scale bar: 100 bp. F) Transgenic *gda-1; xdh-1* animals expressing either wild-type (*Pgda-1::GFP::BsGDA[WT]*) or catalytically inactive (*Pgda-1::GFP::BsGDA[H53A]*) GDA, along with their non-transgenic siblings, were scored for xanthine stone formation. Data points represent biological replicates. Mean and standard deviation are displayed. ns, not significant; *, p<0.05; **, p<0.01; *t* test. Complete details on sample size and individuals scored per biological replicate are provided in **Table S1**.

To determine the site of GDA-1 action, we engineered a wild-type GDA-1 transgene with an N-terminal green fluorescent protein (GFP) fusion driven by the native *gda-1* promoter (*Pgda-1::GFP::GDA-1[WT]*, **Fig. 2B,C**). Transgenic *gda-1; xdh-1* double mutant animals expressing *Pgda-1::GFP::GDA-1* were generated and were analyzed for GFP::GDA-1 expression [26]. GFP::GDA-1 expression was only detected in the *C. elegans* intestine within both the cytoplasmic and nuclear compartments of the cell (**Fig. 2B**). Importantly, the *Pgda-1::GFP::GDA-1* transgene rescued the formation of xanthine stones in *gda-1; xdh-1* double mutants in three independent transgenic lines, strongly suggesting that the intestinal GFP::GDA-1 localization represents the native GDA-1 expression pattern (**Fig. 2D**). We conclude that GDA-1 is expressed and acts in the intestine to limit the formation of xanthine stones.

To determine if *Pgda-1::GFP::GDA-1* rescue was dependent upon a catalytically active deaminase, we engineered a *Pgda-1::GFP::GDA-1[H65A]* construct which altered the essential histidine 65 in the active site of the enzyme. Transgenic *gda-1; xdh-1* double mutant animals expressing *Pgda-1::GFP::GDA-1[H65A]* developed xanthine stones similar to their non-transgenic siblings, demonstrating that H65 was necessary to rescue the xanthine stone phenotype displayed by *gda-1; xdh-1* animals (**Fig. 2D)**. We observe similar GFP::GDA-1 levels between the wild type and H65A variants of the transgene (**Fig. S2A,B**). These results suggest that the xanthine stone accumulation observed in *gda-1; xdh-1* mutant animals is caused by loss of GDA-1 deaminase activity in the *C. elegans* intestine.

To genetically test the hypothesis that GDA-1 is a guanine deaminase, we attempted to rescue *gda-1* loss of function with the well characterized guanine deaminase *Bs*GDA from the bacterium *B. subtilis.* We engineered a transgene expressing *Bs*GDA with an N-terminal GFP fusion expressed by the native *C. elegans gda-1* promoter (*Pgda-1::GFP::BsGDA[WT]*, (**Fig. 2E**) [25, 27]. Transgenic *gda-1; xdh-1* double mutant animals expressing *Pgda-1::GFP::BsGDA* were generated and analyzed for the formation of xanthine stones. GFP::*Bs*GDA was expressed in the intestine and sufficient to rescue the formation of xanthine stones in *gda-1; xdh-1* double mutant animals in three independent transgenic lines (**Fig. 2F).** This rescue was again dependent upon the catalytically essential histidine 53 (**Fig. 2F**). Here we see GFP::*BsGDA* expression in both wild-type and H53A variants of the transgene, although we note that GFP levels appear reduced in the H53A variant (**Fig. S2C,D**). Together, these data support the model that the previously uncharacterized Y48A6B.7 genetic locus encodes a *C. elegans* **<u>g</u>**uanine **<u>d</u>**e**<u>a</u>**minase, which we have named *gda-1* [25, 27].

### *gda-2* encoded a guanine deaminase paralogous to *gda-1*

The *C. elegans* genome encodes a gene (R13A5.10) paralogous to *gda-1* which we name *gda-2*. GDA-1 and GDA-2 display 66% identity at the amino acid sequence level and complete conservation of catalytically essential residues (**Fig. 2A**) [24, 25]. Despite this sequence conservation, *gda-1* and *gda-2* were not functionally redundant with respect to the limiting xanthine stone formation in an *xdh-1* mutant background; *gda-1; xdh-1* double mutant animals displayed xanthine stones even though a wild-type copy of *gda-2* was present in the genome (**Fig. 3A**). Furthermore, *gda-2* loss of function did not enhance the formation of xanthine stones in an *xdh-1* mutant background and did not promote xanthine accumulation (**Fig. 3A,B).**

**Figure 3:**
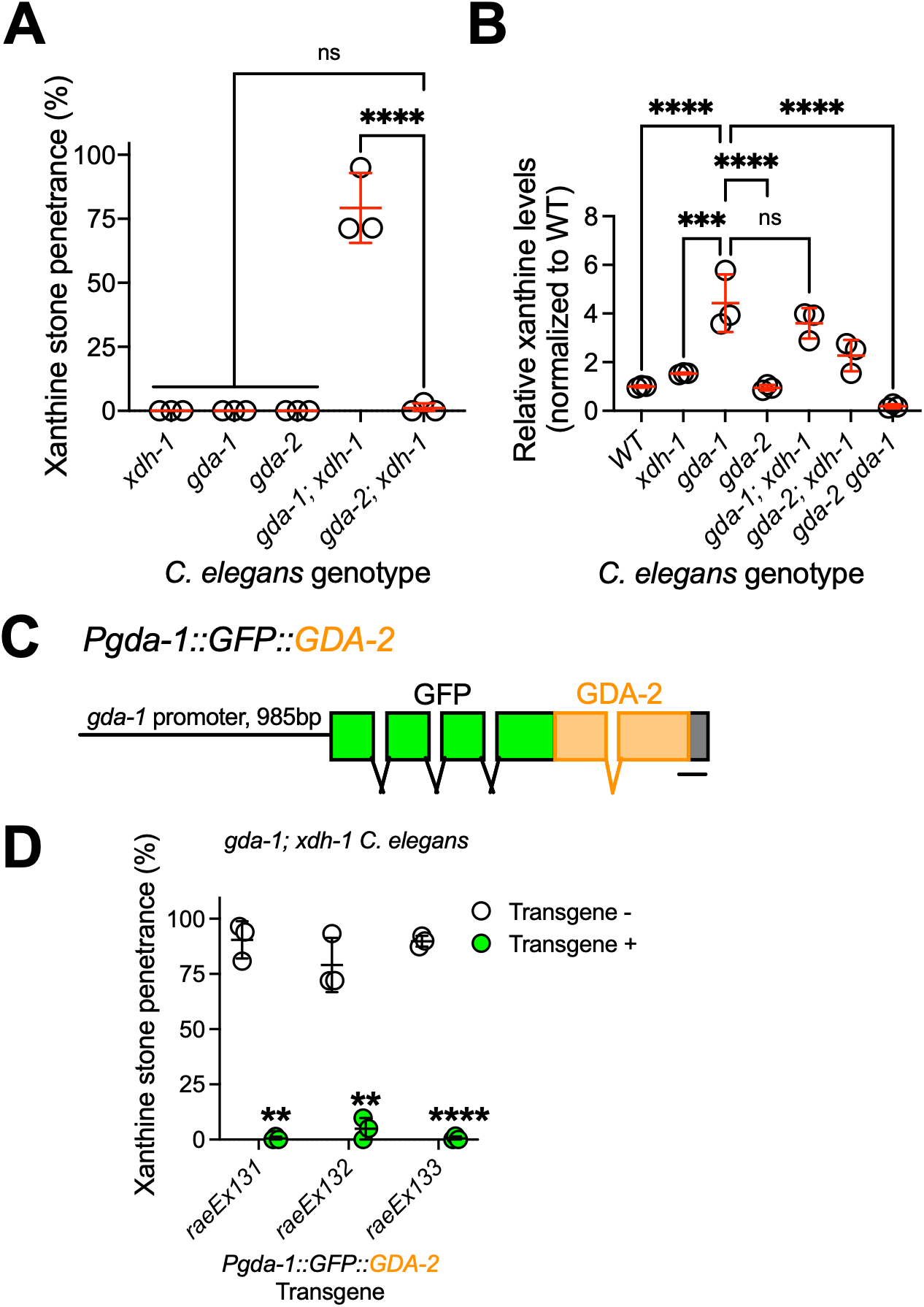
*gda-2* encoded a guanine deaminase paralogous to *gda-1.* A) Xanthine stone formation in various mutant *C. elegans* was assessed. Data points represent biological replicates. Mean and standard deviation are displayed. ns, not significant; ****, p<0.0001; ordinary one-way ANOVA. B) Quantification of xanthine in wild-type and various mutant young adult *C. elegans*. Each data point represents a biological replicate, and values are normalized to the wild type. ns, not significant; ***, p<0.001; ****, p<0.0001; ordinary one-way ANOVA. Note, these data are derived from the same experiment displayed in **Fig. S8**. C) Schematic of the *Pgda-1::GFP::GDA-2* transgene, which encodes wild-type GDA-2 fused to GFP at the N-terminus. Exons are shown as boxes, introns as connecting lines, and the 985 bp *gda-1* promoter as a straight line. Scale bar: 100 bp. D) Transgenic *gda-1; xdh-1* animals expressing wild-type GDA-2 driven by the *gda-1* promoter (*Pgda-1::GFP::GDA-2*), along with their non-transgenic siblings, were scored for xanthine stone formation. Data points represent biological replicates. Mean and standard deviation are displayed. **, p<0.01; ****, p<0.0001; *t* test. Complete details on sample size and individuals scored per biological replicate are provided in **Table S1**.

Given the striking similarity between GDA-1 and GDA-2 amino acid sequences, we hypothesized that *gda-2* encodes an enzyme with guanine deaminase activity, like *gda-1*. To determine if GDA-2 and GDA-1 were functionally interchangeable, we engineered a transgene expressing GDA-2 with an N-terminal GFP tag under the control of the *gda-1* promoter (*Pgda-1::GFP::GDA-2)* (**Fig. 3C**). We sought to test if expressing GDA-2 in the intestine under the control of the *gda-1* promoter was sufficient to rescue *gda-1* loss-of-function. We generated transgenic *gda-1; xdh-1* double mutant *C. elegans* expressing the *Pgda-1::GFP::GDA-2* transgene and assessed the formation of xanthine stones in three independent transgenic lines. *gda-1; xdh-1* mutant animals expressing *Pgda-1::GFP::GDA-2* did not develop xanthine stones, demonstrating functional rescue (**Fig. 3D**). These data demonstrate that *gda-2* is sufficient to rescue *gda-1* mutant loss of function when expressed in the intestine, the site of *gda-1* activity. We suggest that *gda-1* and *gda-2* are functionally interchangeable paralogs that both encode enzymes with guanine deaminase activity.

### *gda-1* and *gda-2* acted redundantly to promote guanine deamination in *C. elegans*

To further test the model that GDA-1 and GDA-2 are guanine deaminases, we measured guanine levels in wild-type, *gda-1, gda-2,* and *gda-2 gda-1* mutant *C. elegans. gda-1* and *gda-2* single mutant *C. elegans* did not accumulate significantly more guanine when compared to the wild type (**Fig. 4A)**. However, *gda-2 gda-1* double mutant animals accumulated 67-fold more guanine than wild type, dramatically increased compared to either *gda-1* or *gda-2* single mutant animals alone (**Fig. 4A)**. Additionally, *gda-2 gda-1* double mutant animals developed dull autofluorescent stones, distinct in fluorescent character from the bright xanthine stones observed in *gda-1; xdh-1* double mutant animals (**Fig. 4B-G**). Given that *gda-2 gda-1* double mutant animals hyperaccumulate guanine and lack xanthine, we propose that these dull autofluorescent stones are likely to be guanine stones (**Fig. 3B and 4A**). Consistent with this model, fluorescent imaging of pure guanine and xanthine revealed similar fluorescent character to the guanine and xanthine stones we observe in our *C. elegans* genetic models (**Fig. S3**). Furthermore, formation of autofluorescent stones in *gda-2 gda-1* double mutant animals required *pnp-1* activity, which catalyzes the production of guanine from guanosine (**Fig. 1A**, **4B**). Taken together, these data are consistent with the model that GDA-1 and GDA-2 are guanine deaminases that act redundantly to limit the accumulation of guanine in *C. elegans*.

**Figure 4:**
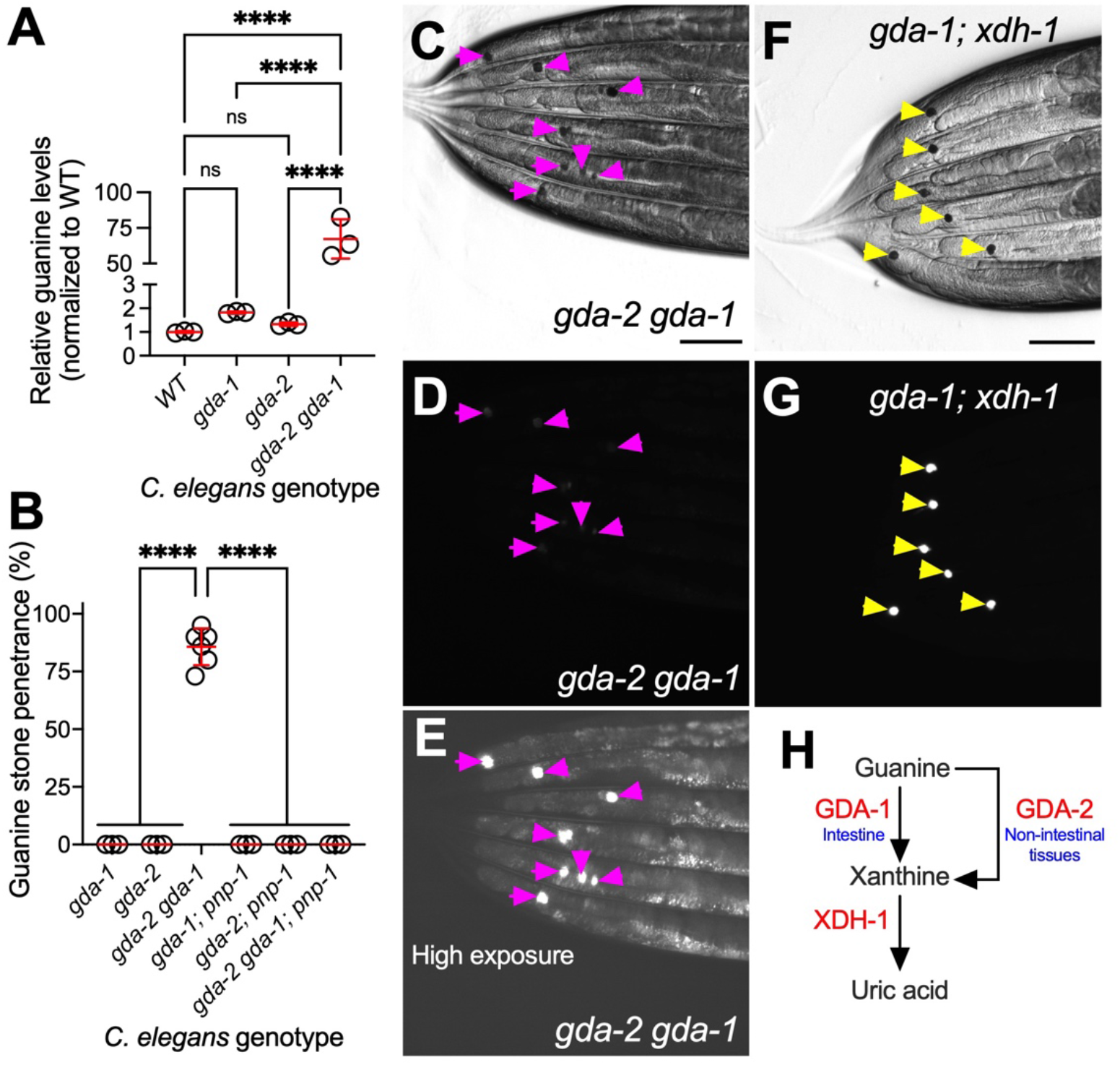
*gda-1* and *gda-2* acted redundantly to limit guanine accumulation. A) Quantification of guanine in wild-type, *gda-1, gda-2,* and *gda-2 gda-1* mutant young adult *C. elegans*. Each data point represents a biological replicate, and values are normalized to the wild type. ns, not significant; ****, p<0.0001; ordinary one-way ANOVA. Note, these data are derived from the same experiment displayed in **Fig. S8**. B) Guanine stone formation in single, double, and triple mutant *C. elegans* was assessed. Data points represent biological replicates. Mean and standard deviation are displayed. ****, p<0.0001; ordinary one-way ANOVA. C-G) Brightfield (C,F) and fluorescence microscopy (D,E,G) images of the posterior section of *gda-2 gda-1* (C-E) and *gda-1; xdh-1* mutant animals (F,G). Pink arrowheads denote weakly autofluorescent guanine stones while yellow arrowheads denote strongly autofluorescent xanthine stones. Note, fluorescence images in D and G were captured at the same exposure and are directly comparable while the image in E was captured with a higher exposure to reveal the weakly autofluorescent guanine stones. Scale bars: 100 μm. H) Model for guanine catabolism facilitated by GDA-1 and GDA-2. During *gda-1* loss of function, GDA-2 activity increases in non-intestinal tissues leading to the hyperaccumulation of xanthine.

Critically, these data shed light on the counterintuitive observation that xanthine, the product of GDA-1-mediated guanine deamination, accumulates during *gda-1* loss of function (**Fig. 1B and 3B**). During *gda-1* loss of function, guanine is deaminated at an increased rate via GDA-2. This is supported by the observation that *gda-2* was necessary for the accumulation of xanthine during *gda-1* loss of function (**Fig. 3B**). In fact, *gda-2 gda-1* double mutant *C. elegans* displayed a substantial reduction in xanthine below that observed in either *gda-1* or *gda-2* single mutant animals (**Fig. 3B**). This result supports the model that *gda-2* activity increases during *gda-1* loss of function resulting in accumulation of xanthine and leading to the formation of xanthine stones when *xdh-1* is inactive (**Fig. 4H)**.

To explore the mechanism of *gda-2* activation during *gda-1* loss of function, we assessed transcriptional and post-transcriptional regulation of *gda-2.* We speculated that increased *gda-2* activity might reflect increased transcription of *gda-2* or other genes involved in *C. elegans* purine metabolism. Yet, *gda-1* inactivation did not promote mRNA accumulation of any of the purine metabolic genes tested via quantitative PCR (**Fig. S4A**). This suggests that *gda-2* activity is not controlled at the level of transcription or mRNA stability. To test the hypothesis that GDA-2 protein levels may be increased during *gda-1* inactivation, we quantified GFP::GDA-2 levels driven by a *Pgda-2::GFP::GDA-2* transgene in both wild-type and *gda-1* mutant *C. elegans* (**Fig. S4B,C**). We observed that GFP::GDA-2 levels were not reliably increased during *gda-1* inactivation using two independent *Pgda-2::GFP::GDA-2* transgenes. Thus, neither *gda-2* mRNA nor protein accumulate during *gda-1* loss of function.

### *gda-1* inactivation reduced fecundity in a *pnp-1* and *gda-2-*dependent manner

We investigated whether purine dyshomeostasis resulting from *gda-1* loss of function compromises *C. elegans* fitness. To test this, we analyzed developmental rate, embryonic hatching, and brood size for wild-type and *gda-1* single mutant *C. elegans.* We observed no defects in larval development or embryonic hatching but found that *gda-1* mutant animals displayed a ∼34% decrease in brood size compared to the wild type (**Fig. S5).** These data demonstrate that *gda-1* promotes reproductive fitness in *C. elegans.* Interestingly, this reduction in brood size was dependent upon functional *pnp-1* and *gda-2*; both *gda-1; pnp-1* and *gda-2 gda-1* double mutant *C. elegans* exhibited higher brood sizes when compared to *gda-1* single mutant animals (**Fig. S5C**). Thus, *pnp-1* and *gda-2* activity limit fecundity when *gda-1* is inactive. These data mirror our results with xanthine accumulation; xanthine accumulates in *gda-1* mutant animals in a *pnp-1* and *gda-2-*dependent manner (**Fig. 1F**, **3B**). This correlation suggests xanthine accumulation may limit fecundity in *C. elegans*, although this model remains to be rigorously tested.

### *gda-2* was expressed and likely acted in multiple tissues to promote guanine deamination

What might explain the maintenance of functionally interchangeable *gda* paralogs in the *C. elegans* genome? We speculated that these paralogs may both provide a fitness advantage through expression in distinct, functionally relevant tissues. *gda-1* was exclusively expressed in the intestine of *C. elegans* (**Fig. 2B**). We hypothesized that *gda-2* would act in non-intestinal tissues to maintain purine homeostasis. To determine the site of action for *gda-2,* we engineered a *gda-2-*expressing transgene with an N-terminal GFP tag driven by the native *gda-2* promoter (*Pgda-2::GFP::GDA-2*). We found that *Pgda-2::GFP::GDA-2* was expressed in multiple sites including neuronal cells and the excretory cell. Notably, we did not detect *Pgda-2::GFP::GDA-2* expression in the intestine (**Fig. S6A**). Importantly, *Pgda-2::GFP::GDA-2* rescued the formation of guanine stones displayed by *gda-2 gda-1* double mutant animals in three independent transgenic strains (**Fig. S6C**). As expected, the *Pgda-2::GFP::GDA-2* transgene did not rescue the formation of xanthine stones displayed by *gda-1; xdh-1* double mutant animals (**Fig. S6B**). We conclude that *gda-1* and *gda-2* are acting in distinct tissues to facilitate guanine deamination.

To identify the primary site of action for GDA-2, we performed tissue specific rescue experiments. GFP::GDA-2 was expressed under the control of the *sng-1* (neurons), *vha-6* (intestine), *myo-3* (muscle), *col-10* (hypodermis), or *sulp-4* (excretory cell) tissue-specific promoters and assessed for ability to rescue the formation of guanine stones displayed by *gda-2 gda-1* double mutant *C. elegans.* Expression of GFP::GDA-2 in the intestine was sufficient to rescue the formation of guanine stones in *gda-2 gda-1* mutant animals (**Fig. S6C**). This result was anticipated given that *gda-1* and *gda-2* are functionally interchangeable and that *gda-1* acts in the intestine. Thus, this result reflects that the *Pvha-6::GFP::GDA-2* transgene is rescuing *gda-1* loss of function. Expression of GFP::GDA-2 in any of the other individual tissues (neurons, muscle, hypodermis, or excretory cell) alone did not provide substantial rescue of guanine stones in *gda-2 gda-1* mutant animals (**Fig. S6C**). All transgenic strains expressed GFP::GDA-2 in the expected tissues (**Fig. S7**). We conclude that *gda-2* is likely acting in multiple tissues to promote guanine deamination.

### Impact of *gda-1, gda-2*, and *xdh-1* on purine metabolite levels in *C. elegans*

To obtain a more comprehensive view of how *gda-1, gda-2,* and *xdh-1* shape purine metabolism in *C. elegans,* we quantified additional purine intermediates (adenosine, guanosine, xanthosine, inosine, hypoxanthine, uric acid, and allantoin) in wild-type and mutant animals (**Fig. S8A**). As expected, uric acid and allantoin were nearly undetectable in all strains lacking *xdh-1.* This is consistent with the essential role of XDH-1 in uric acid production and with uric acid serving as the precursor for allantoin synthesis, presumably via an unidentified *C. elegans* uricase [28]. Loss of both *gda-1* and *gda-2* caused a substantial accumulation of the nucleoside guanosine, mirroring the accumulation of the corresponding base, guanine (**Fig. S8A**). We propose that this buildup reflects reversal of the PNP-1 reaction under condition of guanine excess, shifting metabolic flux back toward the nucleoside. In contrast, xanthine was nearly undetectable in *gda-2 gda-1; xdh-1* triple mutant animals, indicating that all enzymatic routes of xanthine production were genetically eliminated (**Fig. S8A**). Despite this, these triple mutant animals developed bright autofluorescent stones similar to those in *gda-1; xdh-1* double mutants (**Fig. S9**). We speculate that these stones are composed of a distinct, unknown autofluorescent metabolite. Finally, absolute metabolite quantification in wild-type animals revealed that guanine was present at extremely high levels, ∼900-fold more abundant than the next most common metabolite measured, allantoin (**Fig. S8B**). The potential physiological significance of this striking guanine abundance is considered in the Discussion.

### *gda-2* was likely acquired via horizontal gene transfer from Bacillota

There are two classes of guanine deaminases present across the tree of life. The cytidine deaminase-like family, exemplified by *B. subtilis* GDA, and the amidohydrolase family, exemplified by *E. coli* GuaD [25, 29, 30]. These enzymes are distinct in size, domain architecture, and active site composition [25, 29, 31]. To examine the distribution of the two guanine deaminase families across the animal tree of life, we used BLASTp to search the NCBI ClusteredNR sequence database with the amino acid sequences from *B. subtilis* GDA (WP_043857361) or *E. coli* GuaD (CAD6004693.1). We then calculated the mean bit score from the top 50 (where available) hits for representative animal phylum and superphylum (**Fig. 5A**) [24, 32]. Intriguingly, we see the presence of the amidohydrolase family of guanine deaminases (*E. coli* GuaD-like) across the animal tree of life with the notable exception of the nematode lineage (**Fig. 5A**). For example, human guanine deaminase is homologous to the *E. coli* GuaD amidohydrolase [33]. In contrast, we only observe strong evidence of the *B. subtilis-*like guanine deaminase among nematodes (**Fig. 5A**). These data suggest nematodes are unique among animals in their use of a cytidine deaminase-like guanine deaminase.

**Figure 5:**
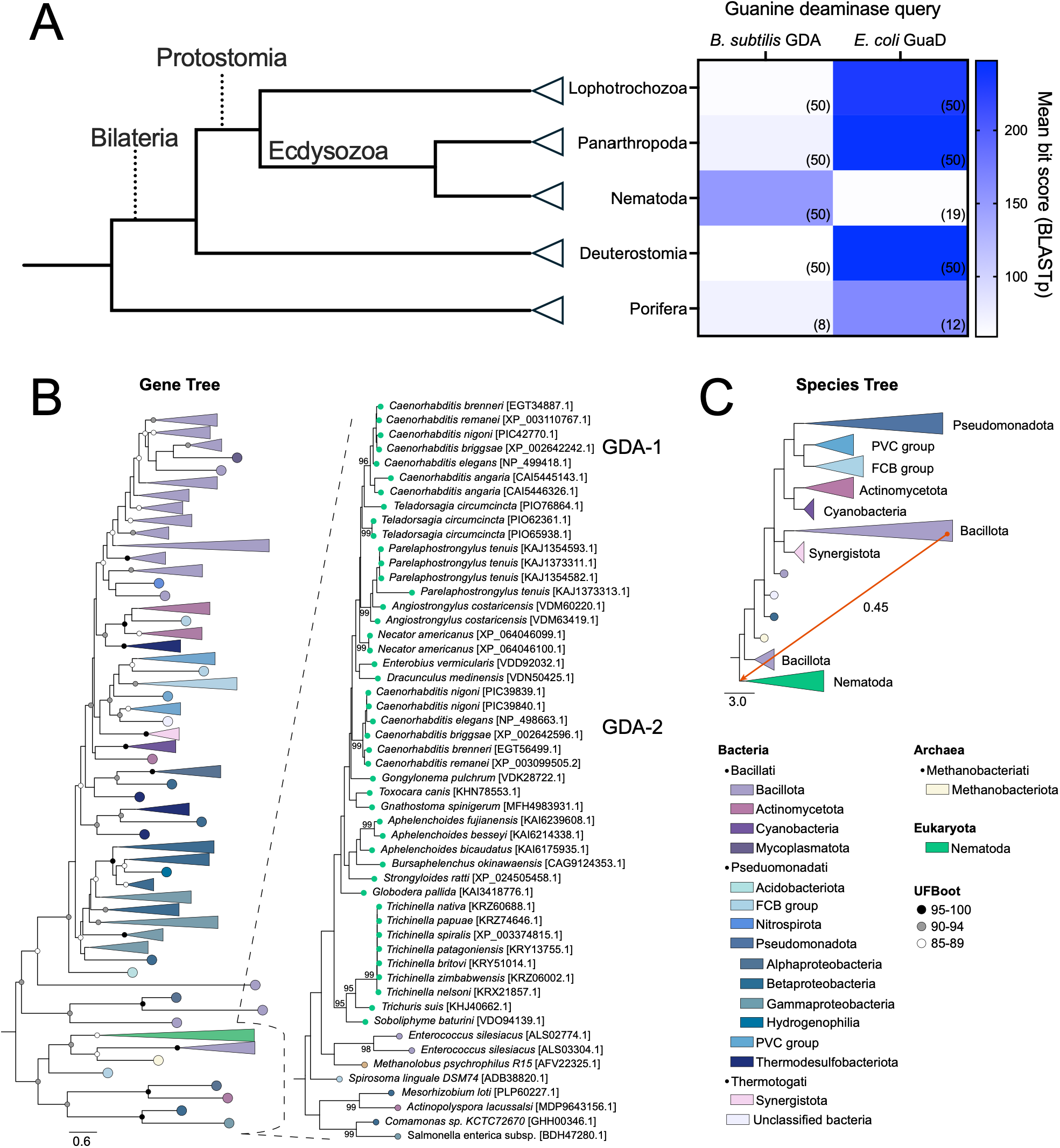
*gda-2* was likely acquired via horizontal gene transfer from Bacillota. A) Cartoon of the animal tree of life (left). The heat map (right) displays the mean BLASTp bit scores using *B. subtilis* GDA and *E. coli* GuaD as queries within each phylum or superphylum. Number of bit scores considered are displayed in parentheses. B) A maximum likelihood gene tree of 257 GDA homologs from 30 nematodes and 207 prokaryotes. The taxa were collapsed and colored at phylum-level. The branch support values (UFBoot2) are indicated by circles: black for 95-100, gray for 90-94, or white for 85-89. A zoomed-in view of the nematode group is shown on right. C) Species tree (Open Tree of Life) with 237 taxa is collapsed and colored based on phylum. A horizontal gene transfer event from Bacillota to the ancestral branch of Nematoda is shown with an inferred transfer rate of 0.45 as calculated by ALE.

Given the intriguing evolutionary pattern of guanine deaminases among animals (**Fig. 5A**), we investigated the evolutionary origins of the nematode GDAs. A BLASTp search against the NCBI ClusteredNR sequence database revealed that the closest matches to *C. elegans* GDA-1 or GDA-2 (highest bit score) were nematode sequences. Outside of nematodes, the most significantly related sequences were all bacterial or archaeal in origin, suggesting a close evolutionary relationship. Based on this pattern of homology, we hypothesized that *gda-1* or *gda-2* entered the nematode lineage via interdomain horizontal gene transfer, rather than by conventional vertical inheritance. This model is consistent with a prior genome-wide analysis of HGT in *C. elegans* that identified *gda-1* or *gda-2* as candidate HGT-derived genes [34]. We sought to test this model given that bioinformatically predicted HGT events do not always withstand individualized scrutiny [35].

To test the HGT model, we identified homologs of GDA-1 and GDA-2 across 30 nematode genomes and 207 prokaryotic genomes and constructed a maximum-likelihood phylogeny. Nematode GDA sequences clustered closely with a subset of Bacillota, supporting a bacterial origin (**Fig. 5B**). Within nematodes, *Caenorhabditis* GDA homologs formed two well-supported clades (UFBoot 96-99), suggesting a duplication event gave rise to the paralogous *gda-1* and *gda-2* within *Caenorhabditis.* In contrast, other nematodes with multiple GDA copies grouped together, indicative of more recent duplication events (**Fig. 5B**). Notably, the *Caenorhabditis* GDA-2 clade branched earlier than GDA-1 suggesting that GDA-2 likely represents the ancestral copy (**Fig. 5B**).

To test for discordance between our maximum-likelihood phylogeny and the tree of life, we performed gene-tree species-tree reconciliation. This analysis inferred a single HGT event from Bacillota into the ancestor of all nematodes (ALE rate=0.45) whereas other potential HGT events were inferred at low rates (ALE rate ∼0.01, **Fig. 5C**). These results support a scenario in which the ancestral nematode gene (*gda-2*) was acquired from Bacillota (possibly a close relative of *Enterococcus silesiacus*), followed by duplication in the *Caenorhabditis* lineage and subsequent diversification. Given the abundance of *E. coli* GuaD-like enzymes in the animal lineage and absence among nematodes (**Fig. 5A)**, we propose that *gda-2* entered the nematode lineage via HGT causing the evolutionary loss of the typical animal guanine deaminase.

Together, our genetic, biochemical, and phylogenetic studies suggest that a guanine deaminase gene from Bacillota entered the nematode lineage via HGT. This gene duplicated, giving rise to *gda-1* and *gda-2*, which subsequently diverged to function in distinct tissues. Our findings trace the evolutionary arc of this transfer and reveal how these acquired genes became integral components of the nematode purine catabolic pathway.

## Discussion

### Purine metabolism is coordinated across multiple tissues in *C. elegans*

Our findings indicate that guanine deamination in *C. elegans* is executed by a distributed metabolic system in which paralogous enzymes act in distinct tissues. GDA-1 functions specifically in the intestine, while GDA-2 likely operates across multiple non-intestinal tissues, revealing that purine catabolism is spatially organized rather than confined in a cell-autonomous manner. This view is consistent with earlier work showing that equilibrative nucleoside transporters ENT-1 and ENT-2 act redundantly to promote development and reproduction [36]. More recent studies demonstrate that these transporters mediate guanosine flux from the intestine to the germline and that combined loss of *ent-1* and *ent-2* depletes germline guanosine, causing sterility [37]. Furthermore, the impact of nucleoside transporters on physiology extends to dietary nutrient availability; *ent-4* encodes an intestinal nucleoside transporter governing dietary nucleoside transport which modulates proteosome function [38]. Collectively, these studies establish that the intestine plays a central role in generating and distributing purine metabolites across tissues.

Within this framework, the distinct consequences of losing GDA-1 versus GDA-2 suggest that guanine, guanosine, xanthine, and related intermediates likely participate in similar inter-tissue transport. The brood-size reduction observed in *gda-1* mutants may therefore reflect not only xanthine accumulation but also altered delivery of guanine-derived nucleosides to metabolically demanding tissues such as the germline. The sensitivity of germline proliferation to somatically supplied purines and the compensatory rewiring of purine biosynthesis observed in ENT-deficient animals further reinforce this possibility [37].

Together, these findings support a model in which purine metabolism in *C. elegans* is inherently multi-tissue, relying on both localized enzymatic processing and transporter-mediated metabolite exchange. As this network is likely more complex than currently appreciated, future work examining how perturbations in transport and enzymatic pathways interact will help clarify the spatial logic of purine homeostasis.

### Loss of intestinal *gda-1* causes unexpected xanthine accumulation

A central feature of our data is that *gda-1* mutants accumulate more xanthine than wild type, even though they lack the major intestinal guanine deaminase. Our proposed model is that loss of intestinal GDA-1 causes guanine to build up and redistribute into distal tissues, where it becomes available to GDA-2 (**Fig. 4H**). While we do not observe regulation of *gda-2* mRNA or protein levels, reaction rates may simply increase with substrate concentration. Thus, elevated regional guanine could increase GDA-2 activity even without changes in *gda-2* expression.

However, substrate availability alone does not explain why xanthine, the product, accumulates. A simple extension of the model is that distal tissues may possess lower XDH-1 activity, producing xanthine faster than it can be cleared. XDH-1 is expressed in the intestine, neurons, and excretory cell—although these tissues have yet to be functionally evaluated for roles in purine catabolism and xanthine stone formation [39]. In this view, wild-type animals deaminate guanine primarily in the intestine, where XDH-1 efficiently catabolizes xanthine; but when GDA-1 is lost, deamination shifts into tissues that are less capable of processing xanthine, allowing it to accumulate.

Finally, because xanthine stones form in the intestinal lumen, xanthine must ultimately move back into the intestine, again indicating that purine intermediates are mobile across tissues. The overall model is therefore that GDA-1 loss redistributes guanine, drives GDA-2 activity in distal tissues, and results in excess xanthine that returns to the intestine where it crystallizes.

### Evolutionary origin and functional diversification of nematode guanine deaminases

Our comparative genomics and phylogenetic analyses reveal that nematodes possess a guanine deaminase lineage with an evolutionary history distinct from that of other animals: nematodes employ a cytidine-deaminase-like guanine deaminase whereas other animals use an amidohydrolase-family guanine deaminase that is notably absent among nematode genomes. We propose that this evolutionary pattern is parsimoniously explained by HGT. However, we cannot exclude the alternative possibility that an ancestral animal possessed both classes of guanine deaminases, followed by repeated loss of the cytidine-deaminase-like guanine deaminase during animal evolution, with the notable exception of the nematode lineage. Supporting the HGT model, gene tree-species tree reconciliation supports a single HGT event from a Bacillota donor into the ancestral nematode lineage, introducing a cytidine-deaminase-like guanine deaminase into nematodes. Given the absence of the animal amidohydrolase-family guanine deaminase in nematodes, we propose that the cytidine-deaminase-like guanine deaminase acquired via HGT has functionally and evolutionarily replaced this enzyme. This evolutionary trajectory raises broader questions about how microbial enzymes become entrenched within core metabolic pathways in animals.

Horizontally transferred genes with clear functional impact are increasingly recognized across metazoans. Coffee berry borer beetles acquired a bacterial mannanase that enables them to access galactomannan, a major polysaccharide in coffee berries that was previously inaccessible [4]. Aphids acquired genes from fungi that promote carotenoid production empowering predator evasion [5]. Nematodes provide several examples of functional HGT. They acquired a bacterial gene that enables L-rhamnose synthesis and strengthens the cuticle against environmental stresses and toxins [6]. They also acquired an algal *cysl* gene that allows them to sense and detoxify amygdalin-derived cyanide, a plant defense compound lethal to most animals [7]. In addition, plant-parasitic nematodes obtained multiple bacterial cell-wall-degrading enzymes that empower them to breach and exploit plant tissues [8]. These examples of HGT suggest clear fitness advantages for the recipient organism.

Our genetic and metabolic data demonstrate that GDA-1 and GDA-2 function as guanine deaminases. Yet the evolutionary advantage of replacing an ancestral amidohydrolase-type enzyme with a horizontally transferred cytidine-deaminase-like guanine deaminase is not obvious and reflects a departure from the classic model of HGT introducing novel biochemical capabilities. Clues to the evolutionary shift may be suggested by our metabolite measurements. We found that wild-type *C. elegans* naturally accumulates high levels of guanine, ∼900-fold more than any other measured purine intermediate. Similar guanine accumulation occurs in organisms that use guanine crystals as metabolic stores or structural materials [40–43]. In fact, the dinoflagellate *Amphidinium carterae* possesses substantial capacity to sequester crystalline guanine, enabling the accumulation of nitrogen reserves sufficient to sustain multiple subsequent generations. These reserves are rapidly built through both direct assimilation of environmental guanine and *de novo* synthesis from diverse nitrogen-containing substrates [42]. We raise the possibility that *C. elegans* also uses guanine as a mobilizable nitrogen reserve. This idea aligns with the species’ natural history, which includes spending a substantial portion of its life in nutrient-limited, diapause-like states [44]. Notably, *B. subtilis* uses guanine as a nitrogen source via a GDA enzyme highly similar with nematode GDA-1 [27]. In fact, we demonstrate that *B. subtilis* GDA is functionally interchangeable with *C. elegans* GDA-1, suggesting that the transferred enzyme may already have been suited for this function upon arrival in the nematode lineage. We propose that the horizontally acquired cytidine-deaminase-like guanine deaminase may have allowed nematodes to exploit guanine as an otherwise inaccessible nitrogen reserve, potentially enhancing survival during nutrient stress. This model remains speculative, and future work will be required to rigorously test it.

Taken together, our analyses support an evolutionary trajectory in which a single bacterial cytidine-deaminase-like guanine deaminase entered the nematode lineage via HGT, followed by loss of the ancestral amidohydrolase-family guanine deaminase. Within *Caenorhabditis*, the horizontally acquired gene duplicated to form *gda-1* and *gda-2*, which then diverged primarily through tissue-specific regulation rather than catalytic specialization. These paralogous enzymes evolved into the sole guanine deaminases driving purine catabolism in nematodes.

## Materials and methods

### General methods and strains

*Caenorhabditis elegans* were maintained at 20°C on nematode growth medium (NGM) plates seeded with *Escherichia coli* OP50, following standard procedures [18]. The wild-type reference strain used was Bristol N2. All mutant and transgenic strains used in this study are listed below. Previously described strains are cited; strains without references were generated in this study.

*Non-transgenic strains:*

N2, wild type [18]

USD869, *xdh-1(ok3234) IV* [17]

USD1044*, gda-1(rae335) III; xdh-1(ok3234) IV*

USD1045*, gda-1(rae336) III; xdh-1(ok3234) IV*

USD1154*, xdh-1(ok3234) IV; cdo-1(mg622) X* [17]

USD1156*, gda-1(rae336) III; cdo-1(mg622) X*

USD1163, *pnp-1(jy121) IV,* outcrossed one time [21]

USD1164, *gda-1(rae336) III; xdh-1(ok3234) IV; cdo-1(mg622) X*

USD1174, *pnp-1(jy121) xdh-1(ok3234) IV* [17]

USD1176, *gda-1(rae336) III; pnp-1(jy121) IV*

USD1188, *gda-1(rae336) III; pnp-1(jy121) xdh-1(ok3234) IV*

USD1178, *gda-2(tm10990) III,* outcrossed four times

USD1179, *gda-2(tm10990) III; xdh-1(ok3234) IV*

USD1227, *gda-2(tm10990) gda-1(rae336) III*

USD1201, *gda-2(tm10990) gda-1(rae336) III; xdh-1(ok3234) IV*

USD1505, *gda-2(tm10990) gda-1(rae336) III; pnp-1(jy121) IV*

USD1508, *gda-2(tm10990) III; pnp-1(jy121) IV*

*Transgenic strains:*

USD1259, *gda-1(rae336) III; xdh-1(ok3234) IV; raeEx126 [Pgda-1::GFP::GDA-1]*

USD1270, *gda-1(rae336) III; xdh-1(ok3234) IV; raeEx135 [Pgda-1::GFP::GDA-1]*

USD1271, *gda-1(rae336) III; xdh-1(ok3234) IV; raeEx136 [Pgda-1::GFP::GDA-1]*

USD1260, *gda-1(rae336) III; xdh-1(ok3234) IV; raeEx127 [Pgda-1::GFP::BsGDA]*

USD1261, *gda-1(rae336) III; xdh-1(ok3234) IV; raeEx128 [Pgda-1::GFP::BsGDA]*

USD1264, *gda-1(rae336) III; xdh-1(ok3234) IV; raeEx130 [Pgda-1::GFP::BsGDA]*

USD1265, *gda-1(rae336) III; xdh-1(ok3234) IV; raeEx131 [Pgda-1::GFP::GDA-2]*

USD1266, *gda-1(rae336) III; xdh-1(ok3234) IV; raeEx132 [Pgda-1::GFP::GDA-2]*

USD1267, *gda-1(rae336) III; xdh-1(ok3234) IV; raeEx133 [Pgda-1::GFP::GDA-2]*

USD1283*, gda-1(rae336)III; xdh-1(ok3234)IV; raeEx140 [Pgda-1::GFP::GDA-1(H65A)]*

USD1284*, gda-1(rae336)III; xdh-1(ok3234)IV; raeEx141 [Pgda-1::GFP::GDA-1(H65A)]*

USD1285*, gda-1(rae336)III; xdh-1(ok3234)IV; raeEx142 [Pgda-1::GFP::GDA-1(H65A)]*

USD1286, *gda-1(rae336)III; xdh-1(ok3234)IV; raeEx143 [Pgda-1::GFP::BsGDA(H53A)]*

USD1287, *gda-1(rae336)III; xdh-1(ok3234)IV; raeEx144 [Pgda-1::GFP::BsGDA(H53A)]*

USD1288, *gda-1(rae336)III; xdh-1(ok3234)IV; raeEx145 [Pgda-1::GFP::BsGDA(H53A)]*

USD1314, *gda-2(tm10990) gda-1(rae336) III; raeEx146 [Pgda-2::GFP::GDA-2]*

USD1315, *gda-2(tm10990) gda-1(rae336) III; raeEx147 [Pgda-2::GFP::GDA-2]*

USD1316, *gda-2(tm10990) gda-1(rae336) III; raeEx148 [Pgda-2::GFP::GDA-2]*

USD1352, *gda-2(tm10990) gda-1(rae336) III; raeEx157 [Psulp-4::GFP::GDA-2]*

USD1353, *gda-2(tm10990) gda-1(rae336) III; raeEx158 [Psulp-4::GFP::GDA-2]*

USD1354, *gda-2(tm10990) gda-1(rae336) III; raeEx159 [Psulp-4::GFP::GDA-2]*

USD1356, *gda-2(tm10990) gda-1(rae336) III; raeEx161 [Pcol-10::GFP::GDA-2]*

USD1357, *gda-2(tm10990) gda-1(rae336) III; raeEx162 [Pcol-10::GFP::GDA-2]*

USD1358*, gda-2(tm10990) gda-1(rae336) III; raeEx163 [Pcol-10::GFP::GDA-2]*

USD1363, *gda-2(tm10990) gda-1(rae336) III; raeEx165 [Psng-1::GFP::GDA-2]*

USD1364, *gda-2(tm10990) gda-1(rae336) III; raeEx166 [Psng-1::GFP::GDA-2]*

USD1365, *gda-2(tm10990) gda-1(rae336) III; raeEx167 [Psng-1::GFP::GDA-2]*

USD1416*, gda-2(tm10990) gda-1(rae336) III; raeEx196 [Pmyo-3::GFP::GDA-2]*

USD1417*, gda-2(tm10990) gda-1(rae336) III; raeEx197 [Pmyo-3::GFP::GDA-2]*

USD1418*, gda-2(tm10990) gda-1(rae336) III; raeEx198 [Pmyo-3::GFP::GDA-2]*

USD1420*, gda-2(tm10990) gda-1(rae336) III; raeEx200 [Pvha-6::GFP::GDA-2]*

USD1421*, gda-2(tm10990) gda-1(rae336) III; raeEx201 [Pvha-6::GFP::GDA-2]*

USD1422*, gda-2(tm10990) gda-1(rae336) III; raeEx203 [Pvha-6::GFP::GDA-2]*

USD1381, *gda-1(rae336) III; xdh-1(ok3234) IV; raeEx180 [Pgda-2::GFP::GDA-2]*

USD1382, *gda-1(rae336) III; xdh-1(ok3234) IV; raeEx181 [Pgda-2::GFP::GDA-2]*

USD1383, *gda-1(rae336) III; xdh-1(ok3234) IV; raeEx182 [Pgda-2::GFP::GDA-2]*

USD1328, *raeEx149 [Pgda-2::GFP::GDA-2]*

USD1329, *raeEx150 [Pgda-2::GFP::GDA-2]*

USD1450*, gda-1(rae336) III; raeEx149 [Pgda-2::GFP::GDA-2]*

USD1451*, gda-1(rae336) III; raeEx150 [Pgda-2::GFP::GDA-2]*

*EMS-derived strains:*

USD972*, xdh-1(ok3234) IV; rae303*

USD973*, xdh-1(ok3234) IV; rae304*

*CRIPSR/Cas9-derived stains:*

USD1035, *gda-1*(*rae335) III*

USD1036, *gda-1(rae336) III*

### Chemical mutagenesis screen and whole genome sequencing

To identify genetic regulators of purine homeostasis, we performed a chemical mutagenesis screen to enhance the penetrance of xanthine stone formation in *xdh-1(ok3234)* mutant *C. elegans* (USD869). Animals were mutagenized using ethyl methanesulfonate (EMS) following established protocols [17, 18].

We report the characterization of two new mutant strains, USD972 and USD973, each harboring EMS-induced lesions (*rae303* and *rae304*, respectively) that enhanced xanthine stone formation in the *xdh-1* mutant background. Both mutations were recessive. When heterozygous, *rae303* and *rae304* caused 0% (n=32) and 1% (n=95) xanthine stone formation in an *xdh-1* mutant background respectively.

To determine whether these mutations affect the same gene, we performed complementation analysis in the *xdh-1(ok3234)* homozygous background. The *rae303* lesion failed to complement *rae304*, with 93% xanthine stone penetrance observed in trans-heterozygotes (n=40), indicating that both mutations disrupt the same genetic locus.

To identify the causative EMS-induced mutations, we performed whole-genome sequencing using established protocols [45]. Genomic DNA was extracted using the Gentra Puregene Tissue Kit (Qiagen), and libraries were prepared with the NEBNext DNA Library Prep Kit (New England Biolabs). Sequencing was conducted on an Illumina NovaSeq platform.

Reads were trimmed and aligned to the *C. elegans* reference genome (WBcel235) using standard workflows on the Galaxy platform [46–48]. Variant calling and annotation were performed to identify deviations from the reference genome and predict functional consequences [49–51].

Both USD972 and USD973 strains shared novel mutations in the gene Y48A6B.*7*, strongly implicating this locus in the enhanced xanthine stone phenotype observed in the USD972 and USD973 mutant strains (**Fig. 1B,C**).

### Quantifying xanthine and guanine stone penetrance

Xanthine and guanine stone formation was quantified using established methods with minor modifications [17]. Briefly, L4-stage animals were seeded onto NGM plates supplemented with 10 µM FUdR to prevent progeny hatching [52]. After 5 days, animals were assessed for the formation of an autofluorescent stone. All data points within each figure panel were collected under identical assay conditions and are directly comparable.

Xanthine stones were identified by the presence of highly autofluorescent puncta that appeared opaque under brightfield microscopy. Guanine stones were identified by the presence of weakly autofluorescent puncta that appeared opaque under brightfield microscopy. Stone penetrance was calculated as the percentage of animals that displayed at least one stone at the time of assessment. Missing or dead animals were excluded from the final analysis.

### Quantifying *C. elegans* life history traits

Developmental rate, hatching efficiency, and brood size were quantified using synchronized populations of *C. elegans*. To determine developmental rate, gravid adult animals were treated with a bleach and sodium hydroxide solution to isolate embryos, which were incubated overnight in M9 buffer to induce hatching and arrest at the L1 stage [53]. These synchronized L1 larvae were cultured 72 hours at 20°C before imaging. Animal length was measured from head to tail using ImageJ software (NIH).

Hatching rate was assessed by collecting eggs from synchronized young adult animals and scoring the number of hatched larvae after approximately 24 hours.

To quantify brood size, individual L4 animals were cloned onto NGM seeded with *E. coli.* Individuals were transferred daily to new plates until the cessation of egg laying. Plates carrying progeny were incubated to allow eggs to hatch, and total brood size was determined by counting the number of hatched progeny. Animals that died or were lost during the assay were excluded from the final analysis.

### CRISPR/Cas9 genome editing

Genome editing of the *gda-1*/Y48A6B.7 locus was performed using established CRISPR/Cas9 protocols [19, 20]. Two guide RNAs (crRNAs; IDT) targeting *gda-1* were designed with the sequences 5′-accccatcacgctgagcaca-3′ and 5′-tatccatgcccaatgtgcat-3′. Ribonucleoprotein complexes consisting of Cas9 protein (IDT) and guide RNAs were assembled and microinjected into the germline of *C. elegans* [19]. Progeny of injected animals were screened for deletions using PCR with the following primers: 5′-ccgtagtaaattgcgtcgaa-3′ and 5′-aaagctcgtcgctgataagg-3′. Two independent deletion alleles, *gda-1(rae335)* and *gda-1(rae336)*, were isolated, made homozygous, and confirmed by Sanger sequencing.

### Microscopy

Fluorescence and low-magnification brightfield images were acquired using a Nikon SMZ25 stereomicroscope equipped with a Hamamatsu ORCA-Flash 4.0 digital camera and NIS-Elements imaging software (Nikon). Fluorescent images of metabolic stones were visualized and imaged using the EGFP BP (FITC/Cy2) HC Filter Set (Nikon). High-magnification differential interference contrast (DIC) and GFP fluorescence images were captured using a Nikon NiE compound microscope, also equipped with a Hamamatsu ORCA-Flash 4.0 digital camera and NIS-Elements software.

All images were processed and analyzed using ImageJ software (NIH). Live *C. elegans* were immobilized using sodium azide prior to imaging. For metabolite imaging, xanthine and guanine (Millipore Sigma) were prepared on microscope slides, covered with coverslips, and sealed with nail polish.

GFP fluorescence was quantified by measuring the average pixel intensity across an entire animal using Image J software (NIH). Background fluorescence was determined from a defined region of interest that did not contain any *C. elegans* and was subtracted from the sample measurements. GFP intensity values were normalized to the wild-type control, which was set to 1.

### Plasmid construction and transgenesis

Plasmid construction was performed using isothermal (Gibson) assembly [54]. For cloning *C. elegans gda-1* and *gda-2,* the loci were amplified from the start codon to the stop codon, including all exons and introns, and assembled with an N-terminal green fluorescent protein (GFP). For cloning *B. subtilis* guanine deaminase, the coding sequence was amplified from start codon to stop codon and assembled with an N-terminal GFP. The *gda-1* promoter was defined as the 985 bp upstream of the *gda-1* start codon. The *gda-2* promoter was defined as the 1,446 bp upstream of the *gda-2* start codon. The *sng-1* neural-specific promoter was defined as the 2,080 bp upstream of the *sng-1* start codon. The *col-10* hypodermal-specific promoter was defined as the 1,129 bp upstream of the *col-10* start codon. The *myo-3* muscle-specific promoter was defined as the 2,168 bp upstream of the *myo-3* start codon. The *sulp-4* excretory cell-specific promoter was defined as the 1,330 bp upstream of the *sulp-4* start codon. The *vha-6* intestine-specific promoter was defined as the 934 bp upstream of the *vha-6* start codon. Site directed mutagenesis was performed on pSB3 and pSB7 to generate pSB9 and pSB10 respectively following manufacturer’s instructions (New England Biolabs). Plasmid sequences were verified by long-read sequencing using Plasmidsaurus, a commercial nanopore-based sequencing service. Plasmids generated for this work are listed below:

pSB3, (*Pgda-1::GFP::GDA-1*)

pSB9, (*Pgda-1::GFP::GDA-1[H65A])*

pSB6, (*Pgda-1::GFP::GDA-2)*

pSB7, (*Pgda-1::GFP::BsGDA)*

pSB10, (*Pgda-1::GFP::BsGDA[H53A])*

pSB13, (*Pgda-2::GFP::GDA-2*)

pSB14, (*Psng-1::GFP::GDA-2*)

pSB15, (*Pcol-10::GFP::GDA-2*)

pSB16, (*Pmyo-3::GFP::GDA-2*)

pSB17, (*Pvha-6::GFP::GDA-2*)

pSB18, (*Psulp-4::GFP::GDA-2*)

Transgenic animals carrying extrachromosomal arrays were generated by injecting the gonad of young adult *C. elegans* with a plasmid of interest (2-20ng/µL), a *Pmyo-2::mCherry* co-injection marker (2ng/µL), and DNA scaffold (78-96 ng/µL KB+ ladder, New England Biolabs) [26, 55].

### mRNA isolation and quantification by qPCR

Wild-type and *gda-1(rae336)* synchronized young adult *C. elegans* were washed in M9 buffer prior to RNA extraction using TRIzol Reagent (Invitrogen), following the manufacturer’s instructions. Total RNA was reverse-transcribed into cDNA using the GoScript Reverse Transcriptase System (Promega).

Quantitative PCR (qPCR) was performed using the CFX96 Real-Time System (Bio-Rad) with SYBR Green Master Mix (Applied Biosystems), according to the manufacturer’s protocols. Relative mRNA levels were calculated using the comparative CT (ΔΔCT) method and normalized to the average expression of *ama-1* mRNA [56]. qPCR primers were:

*ama-1,* 5’-agtgccgagattgaaggaga-3’ and 5’-gtattgcatgttacctttttcaacg-3’

*gda-1,* 5’-attcggtgcggttgtagtt-3’ and 5’-tgggcatggatagcaagatg-3’

*xdh-1:* 5’-gggacgaatgtcggaagaatag-3’ and 5’-cgaatcgagttggtgtcaagta-3’

*gda-2:* 5’-gatgggaaggtaatcggtagtg-3’ and 5’-catggatagcaggaagtgtagag-3’

*hprt-1:* 5’-gtgtgctcgttgtggatgata-3’ and 5’-tccaatccgtagccaacaataa-3’

*pnp-1:* 5’-ggatgcaacgatcccagatt-3’ and 5’-caccctcgtacaacgtcatatc-3’

*adah-1:* 5’-tggtggtccaaaggaagttatt-3’ and 5’-atggaacagcaccagtcatt-3’

*cpin-1:* 5’-catttggaagaacaacccaaaga-3’ and 5’-gctttgagaacaggagcatattg-3’

*gmps-1:* 5’-tgtccagtttcatccagaagtag-3’ and 5’-gacacctccagacaccattac-3’

### Metabolite quantification

Synchronized populations of approximately 5,000 young adult *C. elegans* were cultured under standard conditions. Animals were harvested and washed thoroughly in M9 buffer to remove residual dietary *E. coli*. All subsequent extraction steps took place at 4°C. Worm pellets were flash-frozen in liquid nitrogen and subsequently homogenized with 1.5mm high impact zirconium beads using a BeadBug homogenizer (Benchmark Scientific). *C. elegans* extracts were then centrifuged at 12,000 rpm for five minutes. The supernatant was then filtered through a 10 kDa molecular weight cutoff filter (Pierce), diluted 1:10, and freeze-dried (Harvest Right) to produce the *C. elegans* metabolite extracts.

Metabolites were quantified using isotope dilution mass spectrometry. Freeze-dried *C. elegans* metabolite extracts were resuspended in 50 µL of H₂O containing 0.0075% (v/v) formic acid and the appropriate isotope-labeled internal standards. Samples were vortexed for 20 seconds and incubated at 30°C with shaking at 1000 rpm for 30 minutes. After centrifugation at 50,000 x g for 30 minutes, the supernatant was transferred to mass spectrometry vials.

Quantification of nucleosides, nucleobases, uric acid, and allantoin was performed using an Agilent 1200 HPLC system coupled to an Agilent 6470C triple quadrupole mass spectrometer. Chromatographic separation was achieved using a Polaris 5 C18-A column (50 × 4.6 mm; Agilent Technologies) operated at 30°C with a flow rate of 0.650 mL/min. Mobile phase A consisted of: 75 µL/L formic acid in 100% H₂O and mobile phase B consisted of 75 µL/L formic acid in 100% methanol (MeOH). The injection volume was 10 µL. Analyses were conducted in positive ion mode for nucleosides and nucleobases, and in negative ion mode for uric acid and allantoin, using multiple-reaction monitoring (MRM).

### Evolutionary analyses of *C. elegans* guanine deaminases

To approximate the distribution of the two classes of guanine deaminases across the tree of life, we used *B. subtilis* GDA (WP_043857361) or *E. coli* GuaD (CAD6004693.1) as BLASTp queries. For each guanine deaminase query, a BLASTp search against the ClusteredNR database (NIH) of various animal phylum and superphylum was performed. For each BLASTp search, the top 50 bit scores were collected. When 50 significant bit scores were not returned, all significant bit scores were considered.

To evaluate potential horizontal gene transfer events, a local nematode database was built using 37 nematode genomes obtained from WormBase [57]. Using *C. elegans* proteins GDA-1 (NP_499418.1) and GDA-2 (NP_498663.1) as query, 52 nematode homologs were curated using BLASTp [58]. These nematode sequences were aligned using MAFFT v7.49 to generate an HMM profile with HMMER v3.3.2 (*hmmbuild*) [59, 60]. To identify prokaryotic homologs, the same *C. elegans* query proteins were searched via BLASTp against a local GTDB-derived database containing prokaryotic genomes with >95% completeness and <5% contamination. The BLAST hits were further filtered using the nematode-based HMM profile (*hmmsearch*) to increase specificity [59]. The resulting prokaryotic sequences were clustered at 70% identity using CD-HIT [61], resulting a dataset of 722 prokaryotic proteins.

To obtain a species tree for gene/species tree reconciliation, the organisms in the protein dataset were searched against Open Tree of Life [62], identifying 237 matching taxa which were retrieved for species tree. GDA protein sequences (n=257) that belong to 237 taxa were aligned using MAFFT v7.49 and a maximum-likelihood gene tree was constructed using IQTREE v2.0.3 by selecting LG+R10 as evolutionary model using ModelFinder [60, 63, 64]. Gene/species tree reconciliation was then performed using Amalgamated Likelihood Estimation (ALE) calculating the rates of horizontal gene transfer events [65].

### Declaration of generative AI and AI-assisted technologies in the writing process

During the preparation of this work, the author(s) used Microsoft Copilot to improve the clarity and readability of the text. All content was reviewed and edited by the author(s), who take full responsibility for the final version of the manuscript.

## Acknowledgments

Some *C. elegans* strains were provided by the CGC, which is funded by the NIH Office of Research Infrastructure Programs (P40 OD010440). *gda-2(tm10990)* was provided by the National Bio-Resource Project, which is funded by the Japanese government. This work was funded by the National Institutes of Health (NIH) National Institute of General Medical Sciences (R35GM146871 to KW), an NSF REU (1756912 to GH), the UW-Madison Alumni Foundation Hiroshi-Sugiyama Fund (EF), the UW-Madison Bacteriology Department Peterson Fellowship (EF), and a NASA Astrobiology Program Grant (NNH23ZDA001N-EXO to BK). The funders had no role in study design, data collection and analysis, decision to publish, or preparation of the manuscript. We would also like to thank the Center for High Throughput Computing (CHTC) at the University of Wisconsin-Madison for providing computing resources.

## Supplemental Figures

**Figure S1:**
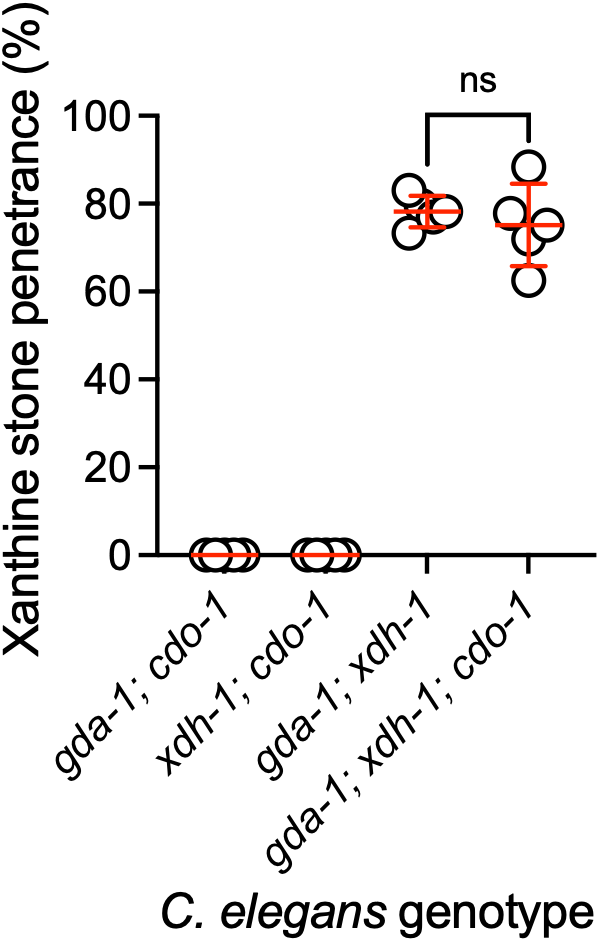
*cdo-1* was not required for xanthine stone formation in *gda-1; xdh-1* mutant animals. Xanthine stone formation in various mutant *C. elegans* was assessed. Data points represent biological replicates. Mean and standard deviation are displayed. ns, not significant; *t* test.

**Figure S2:**
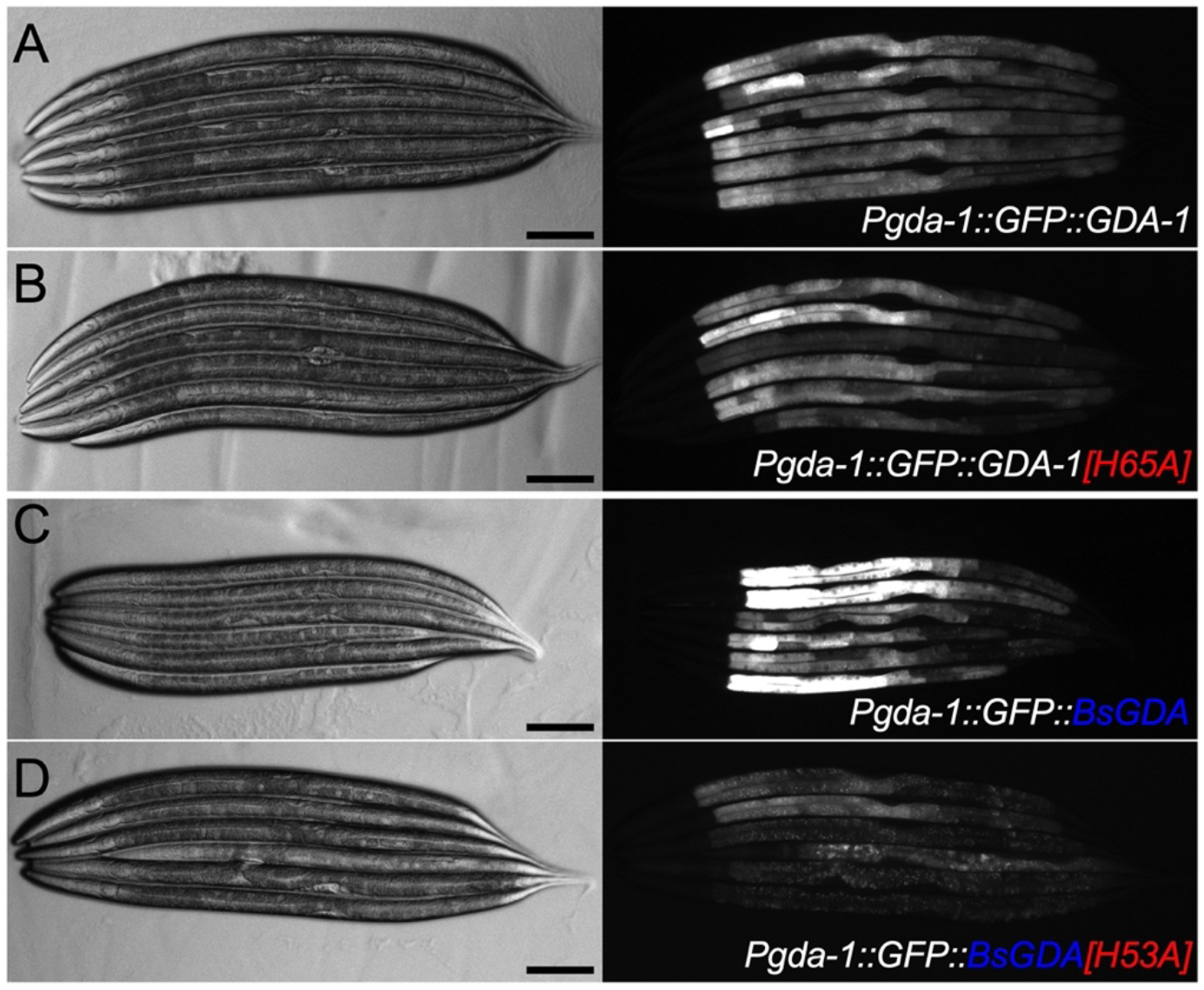
Expression of various *C. elegans* transgenes. Representative brightfield (left) and fluorescence (right) images of *gda-1; xdh-1* double mutant *C. elegans* expressing A) *Pgda-1::GFP::GDA-1,* B) *Pgda-1::GFP::GDA-1[H65A]*, C) *Pgda-1::GFP::BsGDA,* or D) *Pgda-1::GFP::BsGDA[H53A]* transgenes. Animals were imaged at the L4 stage of development. Note, fluorescence images in A-C are directly comparable (20ms exposure) while fluorescence image in D was taken at a higher 50ms exposure to visualize GFP. Scale bars: 100 μm.

**Figure S3:**
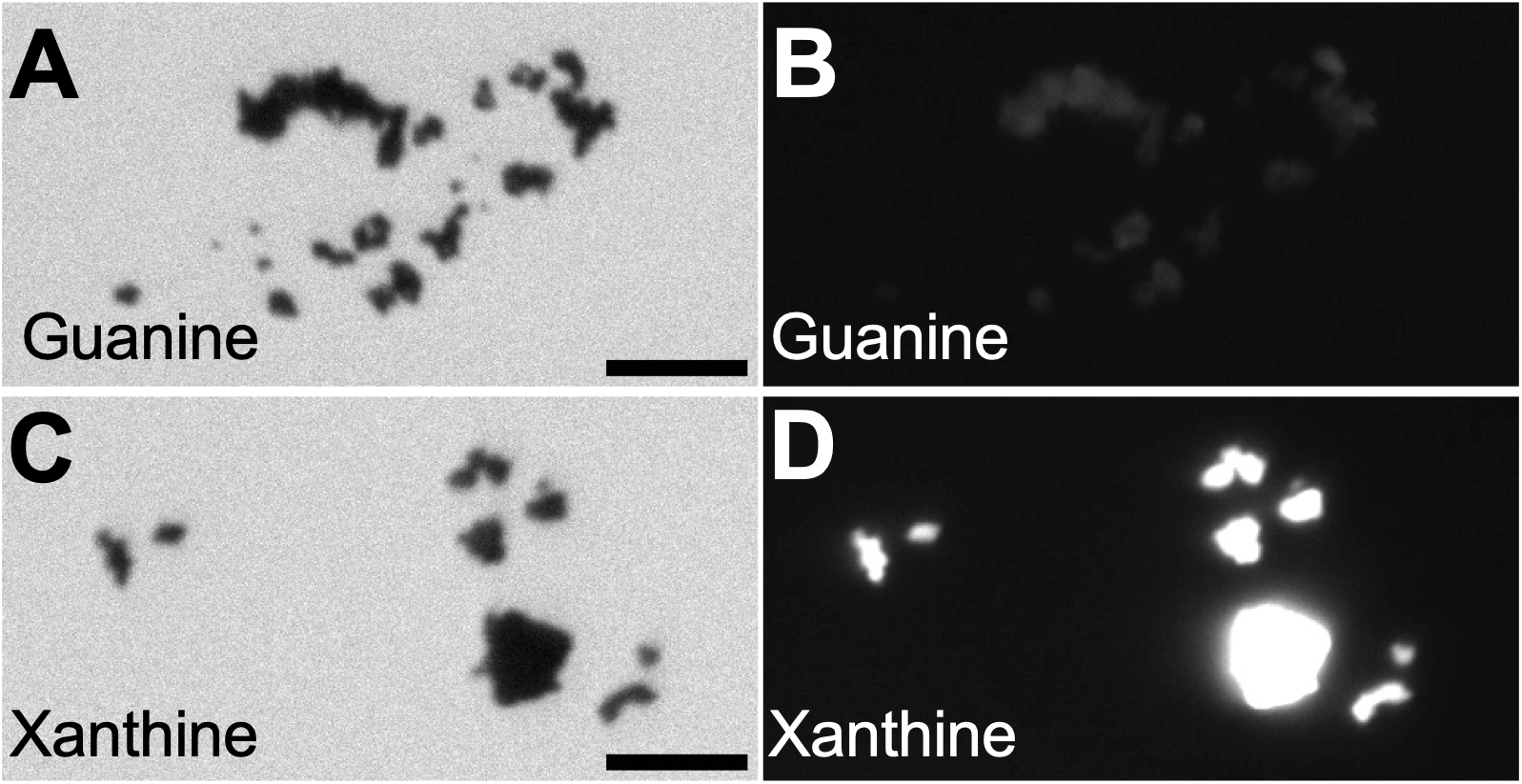
Xanthine and guanine were autofluorescent metabolites. Brightfield (A,C) and fluorescence (B,D) images of guanine (A,B) and xanthine (C,D). Scale bars: 25 μm.

**Figure S4:**
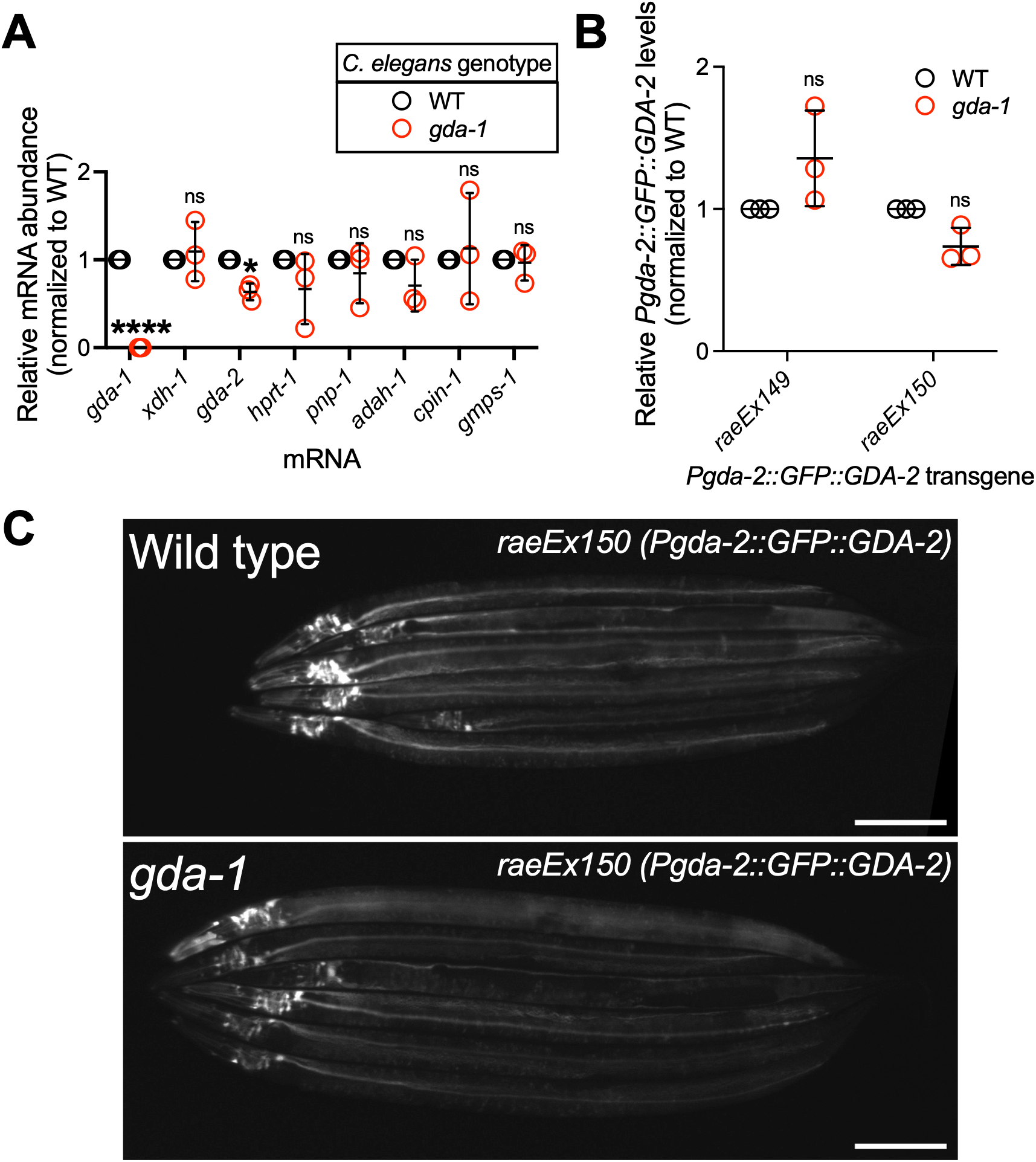
*gda-1* loss of function did not increase *gda-2* mRNA or protein levels in *C. elegans.* A) Total RNA was isolated from wild-type and *gda-1(rae336)* animals at the young adult stage. Transcript levels were quantified by qPCR and analyzed using the delta-delta C_T_ method. All values were normalized to *ama-1* expression, and relative abundance for each transcript was set to 1 in wild type. Individual data points represent biological replicates; mean and standard deviation are displayed. ns, not significant; *, p<0.05; ****, p<0.0001; *t* test. B) Quantification of GFP expression from two distinct extrachromosomal arrays carrying the *Pgda-2::GFP::GDA-2* transgene. Wild-type and *gda-1(rae336)* animals expressing *Pgda-2::GFP::GDA-2* were imaged at the L4 stage of development. Individual datapoints are biological replicates; mean and standard deviation are displayed. ns, not significant; *t* test. Complete details on sample size and individuals scored per biological replicate are provided in **Table S1**. C) Representative fluorescence images of the *Pgda-2::GFP::GDA-2* (*raeEx150*) transgene in wild-type and *gda-1(rae336)* mutant *C. elegans* at the L4 stage. Scale bars: 100 μm.

**Figure S5:**
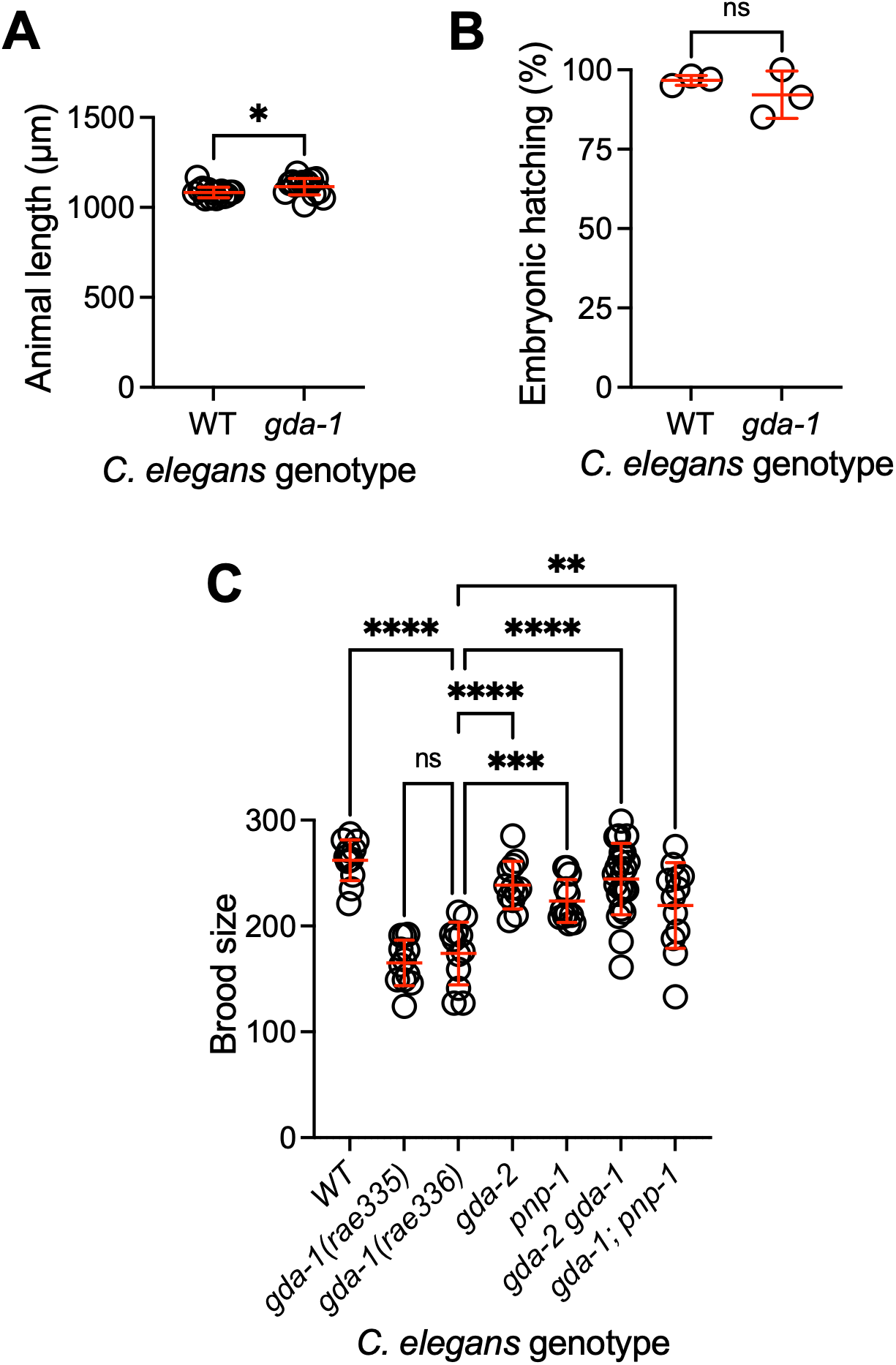
*gda-1* promoted *C. elegans* fecundity. A) Wild-type and *gda-1(rae336)* mutant *C. elegans* were synchronized at the L1 larval stage and cultured for 72 hours. Animal length was measured for each individual. Data are presented as individual points with mean and standard deviation. Sample size: 15 animals per genotype. *, p<0.05; *t* test. B) The hatching rate of newly laid wild-type and *gda-1(rae336)* mutant *C. elegans* embryos was measured. Each data point represents a biological replicate. Mean and standard deviation are shown. ns, not significant; *t* test. C) Brood size of wild-type and mutant *C. elegans* was measured. Each data point represents a biological replicate. Mean and standard deviation are shown. **, p<0.01; ***, p<0.001; ****, p<0.0001; ordinary one-way ANOVA. Complete details on sample size and individuals scored per biological replicate are provided in **Table S1**.

**Figure S6:**
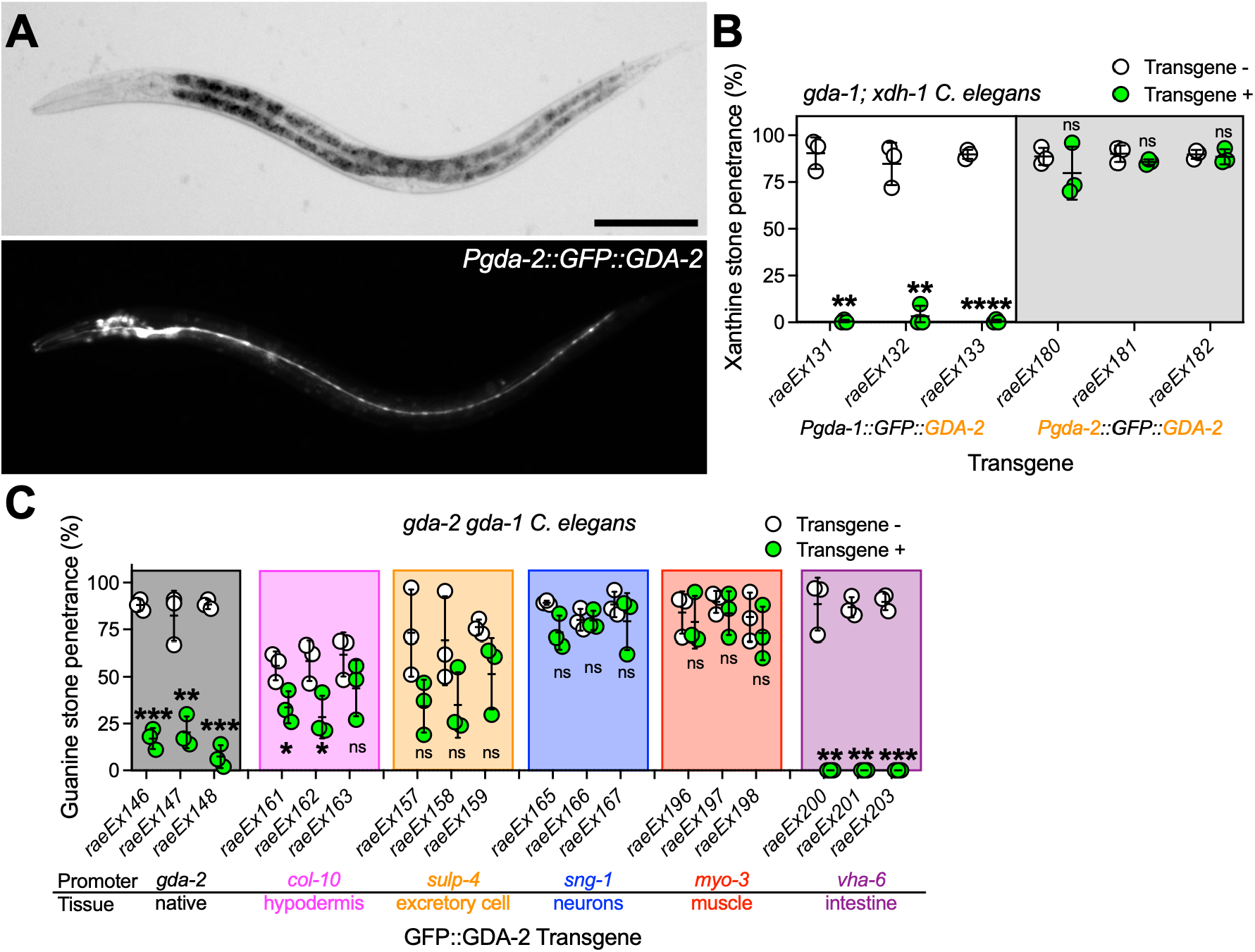
GDA-2 was expressed in multiple tissues. A) Brightfield (top) and fluorescence (bottom) images of *gda-2 gda-1* double mutant *C. elegans* expressing the *Pgda-2::GFP::GDA-2* transgene. Animals were imaged at the L4 stage of development. Scale bar: 100 μm. B) Transgenic *gda-1; xdh-1* animals expressing *Pgda-1::GFP::GDA-2* or *Pgda-2::GFP::GDA-2*, along with their non-transgenic siblings, were scored for xanthine stone formation. Data points represent biological replicates. Mean and standard deviation are displayed. ns, not significant; **, p<0.01; ****, p<0.0001; *t* test. C) Guanine stone formation was assessed in transgenic *gda-2 gda-1 C. elegans* expressing *GFP::GDA-2* driven by the native promoter (*gda-2)* and promoters specific to hypodermis (*col-10*), excretory cell (*sulp-4*), neurons (*sng-1*), muscle (*myo-3*), or intestine (*vha-6*). Three independent extrachromosomal arrays were assessed for each tissue of interest. Data points represent biological replicates. ns, not significant; *, p<0.05; **, p<0.01; ***, p<0.001; *t* test. Mean and standard deviation are displayed. Complete details on sample size and individuals scored per biological replicate are provided in **Table S1**.

**Figure S7:**
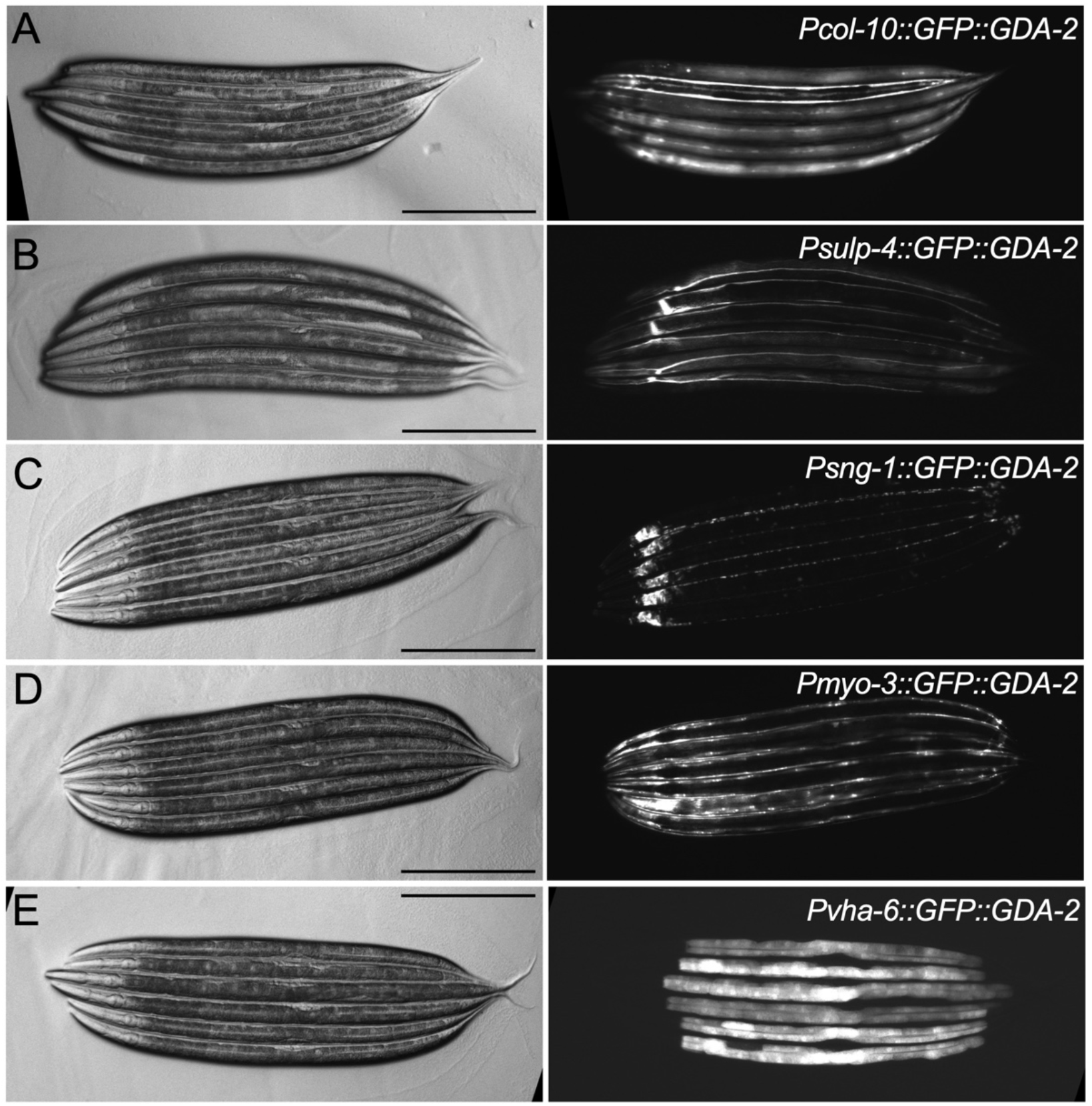
Expression of various *C. elegans* transgenes. Representative brightfield (left) and fluorescence (right) images of *gda-2 gda-1* double mutant *C. elegans* expressing *GFP::GDA-2* transgene driven by the A) hypodermal *col-10,* B) excretory cell *sulp-4*, C) neuronal *sng-1,* D) muscle *myo-3,* or E) intestinal *vha-6* promoters. Animals were imaged at the L4 stage of development. Scale bars: 100 μm.

**Figure S8:**
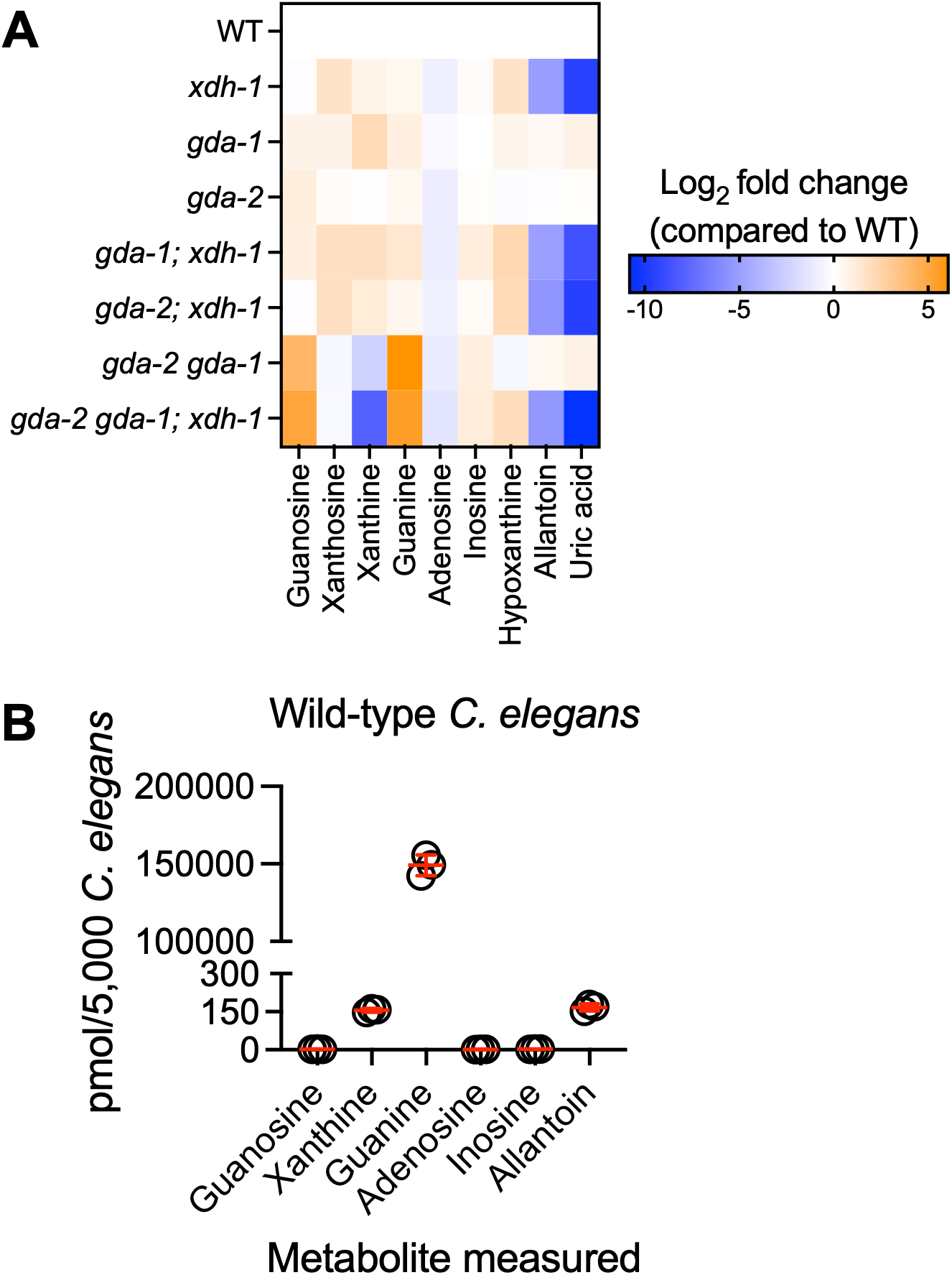
Analysis of purine metabolites in wild-type and mutant *C. elegans.* A) Heat map displaying the relative (log_2_ fold change) amounts of purine metabolites for wild-type and various mutant *C. elegans.* Values are normalized to the wild type. Each cell represents the average of three biological replicates. Raw data are provided in **Table S1**. B) Quantification of purine metabolites from extracts of wild-type young adult *C. elegans*. Individual datapoints are biological replicates; mean and standard deviation are displayed.

**Figure S9:**
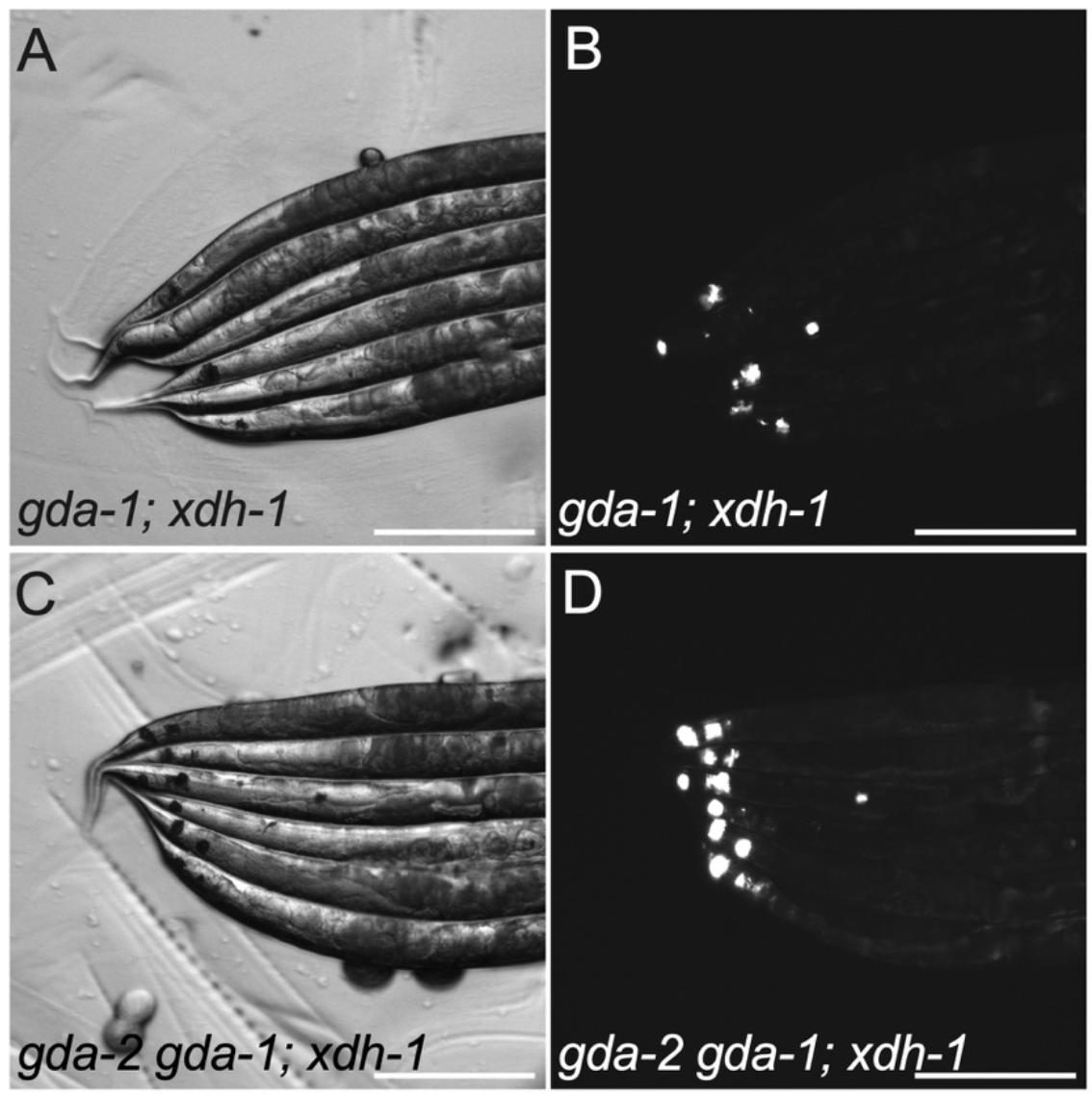
Bright autofluorescent stones displayed by *gda-2 gda-1; xdh-1* mutant *C. elegans.* Brightfield (A,C) and fluorescence (B,D) images are displayed of the posterior of *gda-1; xdh-1* (A,B) or *gda-2 gda-1; xdh-1* (C,D) mutant *C. elegans* at day three of adulthood. Scale bars: 250 μm.

**Table S1: Raw data and information regarding sample sizes and biological replicates.**

